# Cortical ripples during NREM sleep and waking in humans

**DOI:** 10.1101/2021.05.11.443637

**Authors:** Charles W. Dickey, Ilya A. Verzhbinsky, Xi Jiang, Burke Q. Rosen, Sophie Kajfez, Emad N. Eskandar, Jorge Gonzalez-Martinez, Sydney S. Cash, Eric Halgren

## Abstract

Hippocampal ripples index the reconstruction of spatiotemporal neuronal firing patterns essential for the consolidation of memories in the cortex during non-rapid eye movement sleep (NREM). Recently, cortical ripples in humans have been shown to enfold the replay of neuron firing patterns during cued recall. Here, using intracranial recordings from 18 patients (12 female), we show that cortical ripples also occur during NREM in humans, with similar density, oscillation frequency (∼90 Hz), duration, and amplitude to waking. Ripples occurred in all cortical regions with similar characteristics, unrelated to putative hippocampal connectivity, and were less dense and robust in higher association areas. Putative pyramidal and interneuron spiking phase-locked to cortical ripples during NREM, with phase delays consistent with ripple generation through pyramidal-interneuron feedback. Cortical ripples were smaller in amplitude than hippocampal ripples, but were similar in density, frequency, and duration. Cortical ripples during NREM typically occurred just prior to the upstate peak, often during spindles. Upstates and spindles have previously been associated with memory consolidation, and we found that cortical ripples grouped co-firing between units within the window of spike-timing-dependent plasticity. Thus, human NREM cortical ripples are: ubiquitous and stereotyped with a tightly focused oscillation frequency; similar to hippocampal ripples; associated with upstates and spindles; and associated with unit co-firing. These properties are consistent with cortical ripples possibly contributing to memory consolidation and other functions during NREM in humans.

**Significance Statement:** In rodents, hippocampal ripples organize replay during sleep to promote memory consolidation in the cortex, where ripples also occur. However, evidence for cortical ripples in human sleep is limited, and their anatomical distribution and physiological properties are unexplored. Here, using human intracranial recordings, we demonstrate that ripples occur throughout the cortex during waking and sleep with highly stereotyped characteristics. During sleep, cortical ripples tend to occur during spindles on the down-to-upstate transition, and thus participate in a sequence of sleep waves that is important for consolidation. Furthermore, cortical ripples organize single unit spiking with timing optimal to facilitate plasticity. Therefore, cortical ripples in humans possess essential physiological properties to support memory and other cognitive functions.

## Introduction

Hippocampal ripples have been extensively studied in rodents during non-rapid eye movement sleep (NREM), when they mark the replay of events from the prior waking period, and are critical for memory consolidation in the cortex (Wilson and McNaughton, 1994; Ego-Stengel and Wilson, 2009; Girardeau et al., 2009; Buzsaki, 2015; Maingret et al., 2016). They are associated with cortical replay (Ji and Wilson, 2007; Peyrache et al., 2009; Johnson et al., 2010), and with cortical sleep waves (spindles, downstates, upstates) (Siapas and Wilson, 1998), a relationship crucial for consolidation (Latchoumane et al., 2017). Rat hippocampal ripples comprise a ∼140 Hz oscillation riding on the peak of a ∼70 ms duration sharpwave, followed by a slower local potential (Buzsaki, 2015). Human hippocampal sharpwave-ripples also occur during NREM with similar temporal relationships to cortical spindles and down-to-upstates, and similar hippocampal topography, but with a median frequency of 80-90 Hz (Staresina et al., 2015; Jiang et al., 2019a; Jiang et al., 2019b; Jiang et al., 2019c).

Recently, ripples were found in rat association cortex but not primary sensory or motor cortices during sleep, with increased coupling to hippocampal ripples in sleep following learning (Khodagholy et al., 2017). An earlier study reported ripples in waking and sleeping cat cortex, especially during NREM (Grenier et al., 2001). In humans, cortical ripples during waking were more frequently found in lateral temporal than rolandic cortex, and coupled to parahippocampal gyrus ripples more often prior to correct paired-associates recall (Vaz et al., 2019). Lateral temporal units fired in-phase with the local waking ripples in patterns previously observed during learning (Vaz et al., 2020). Waking hippocampal ripples were also associated with cortical activity patterns selective for faces and buildings during free recall (Norman et al., 2019). Evidence for cortical ripples during NREM in humans is limited, but a previous study indicated that they may be suppressed during and increased following the cortical downstate (von Ellenrieder et al., 2016).

Thus, there is an emerging appreciation that, in humans and rodents, hippocampal and cortical ripples play an important role in memory during both sleep and waking. However, many fundamental questions remain unresolved. The basic characteristics of ripples have not been compared between the cortex and hippocampus, or between sleep and waking, so it is unclear how ripples may differ between their putative roles of supporting consolidation vs. recall, or indeed if they represent the same phenomenon. Knowledge of the distribution of ripples across different cortical areas during waking is limited and during sleep is essentially absent. The relations between cortical ripples and local sleep spindles, downstates, and upstates have not been determined. Such relations could support a role of cortical ripples in consolidation, as would increased co-firing between neurons within the window of spike-timing-dependent plasticity (STDP). Furthermore, the relationships of human cortical pyramidal and inhibitory cell-firing to ripples and to each other, important for understanding ripple generation, have not been determined.

Here, using intracranial stereoelectroencephalography (SEEG) recordings, we show that cortical ripples are generated during NREM in humans, and we provide the first comprehensive characterization of cortical ripples. Ripples with a stereotyped, tightly focal oscillation frequency of ∼90 Hz and duration of ∼70 ms were ubiquitous throughout the cortex during waking and NREM, although slightly less dense and robust in association areas, and with no relationship to putative hippocampal connectivity. We found that cortical ripples are similar to hippocampal in oscillation frequency, density, and duration. Cortical ripples in NREM coupled strongly to down-to-upstates, and less often to spindles, consistent with a possible role in memory replay. Using single-unit recordings from cortical microarrays, we identify the probable generating circuits of cortical ripples and show that units co-fire during ripples at short delays that are optimal for STDP. Thus, human cortical ripples during NREM have the necessary physiological properties to facilitate replay-guided plasticity. However, the ubiquity and stereotypy of human ripples across structures and states are also consistent with a more general functional role.

## Results

### Ripples are ubiquitous across states and structures with a characteristic and focal frequency (∼90 Hz) and duration (∼70 ms)

Ripples were detected in intracranial cortical and hippocampal SEEG recordings from 17 patients undergoing monitoring for seizure focus localization (Table 1). Bipolar transcortical derivations ensured measurement of locally-generated LFPs. Ripples were detected exclusively from non-lesional, non-epileptogenic regions and were required to have at least 3 cycles of increased 70-100 Hz amplitude without contamination by epileptiform activity or artifacts (Figure 1). Recording epochs and channels with possible contamination by epileptiform activity were rigorously rejected. Ripples were found during both waking and NREM in all cortical areas sampled (Figure 2A-B,E-J, Table 2) as well as the hippocampus (Figure 2C-D,K-L, Table 2).

**Table 1.** Stereoelectroencephalography patient demographics and data characteristics. Channels are the bipolar derivations included in the analyses. δ NREM / δ Waking reports the ratio of the mean of the delta (0.5-2 Hz) analytic amplitude means across cortical channels during the NREM vs. waking epochs analyzed.

| Patient | Age | Sex | Handedness | No. Cort. Ch. | No. Hipp. Ch. | NREM Dur. (hr) | Waking Dur. (hr) | $\delta$ NREM / $\delta$ Waking |
| --- | --- | --- | --- | --- | --- | --- | --- | --- |
| S1 | 20 | M | R | 18 (L) | 1 (L) | 6.1 | 49.8 | 3.98 |
| S2 | 58 | F | R | 22 (L) | 2 (L) | 23.7 | 23.8 | 2.35 |
| S3 | 42 | M | L | 16 (3 L) | 3 (2 L) | 11.4 | 19.3 | 3.39 |
| S4 | 18 | F | L | 15 (R) | 2 (R) | 2.7 | 2.2 | 3.02 |
| S5 | 20 | F | R | 18 (7 L) | 2 (R) | 5.6 | 19.3 | 2.93 |
| S6 | 22 | M | LR | 17 (L) | 1 (L) | 6.3 | 59.9 | 5.45 |
| S7 | 30 | F | R | 13 (2 L) | 1 (R) | 20.5 | 76.7 | 4.46 |
| S8 | 43 | F | R | 12 (L) | 2 (L) | 8.1 | 32.6 | 5.43 |
| S9 | 16 | M | R | 16 (4 L) | 1 (R) | 16.3 | 4.8 | 2.50 |
| S10 | 32 | F | R | 29 (3 L) | 3 (1 L) | 11.2 | 27.6 | 3.11 |
| S11 | 21 | F | L | 8 (4 L) | 3 (2 L) | 16.0 | 52.6 | 3.04 |
| S12 | 21 | F | R | 14 (13 L) | 1 (L) | 26.2 | 37.2 | 2.92 |
| S13 | 29 | F | R | 15 (6 L) | 2 (1 L) | 8.6 | 24.6 | 3.75 |
| S14 | 41 | F | R | 18 (R) | 1 (R) | 11.9 | 63.3 | 4.20 |
| S15 | 24 | M | R | 21 (9 L) | 1 (L) | 11.8 | 11.6 | 3.27 |
| S16 | 31 | F | R | 15 (7 L) | 1 (L) | 28.1 | 39.4 | 2.55 |
| S17 | 21 | M | R | 6 (L) | 1 (L) | 11.3 | 18.9 | 1.41 |

**Table 2.**
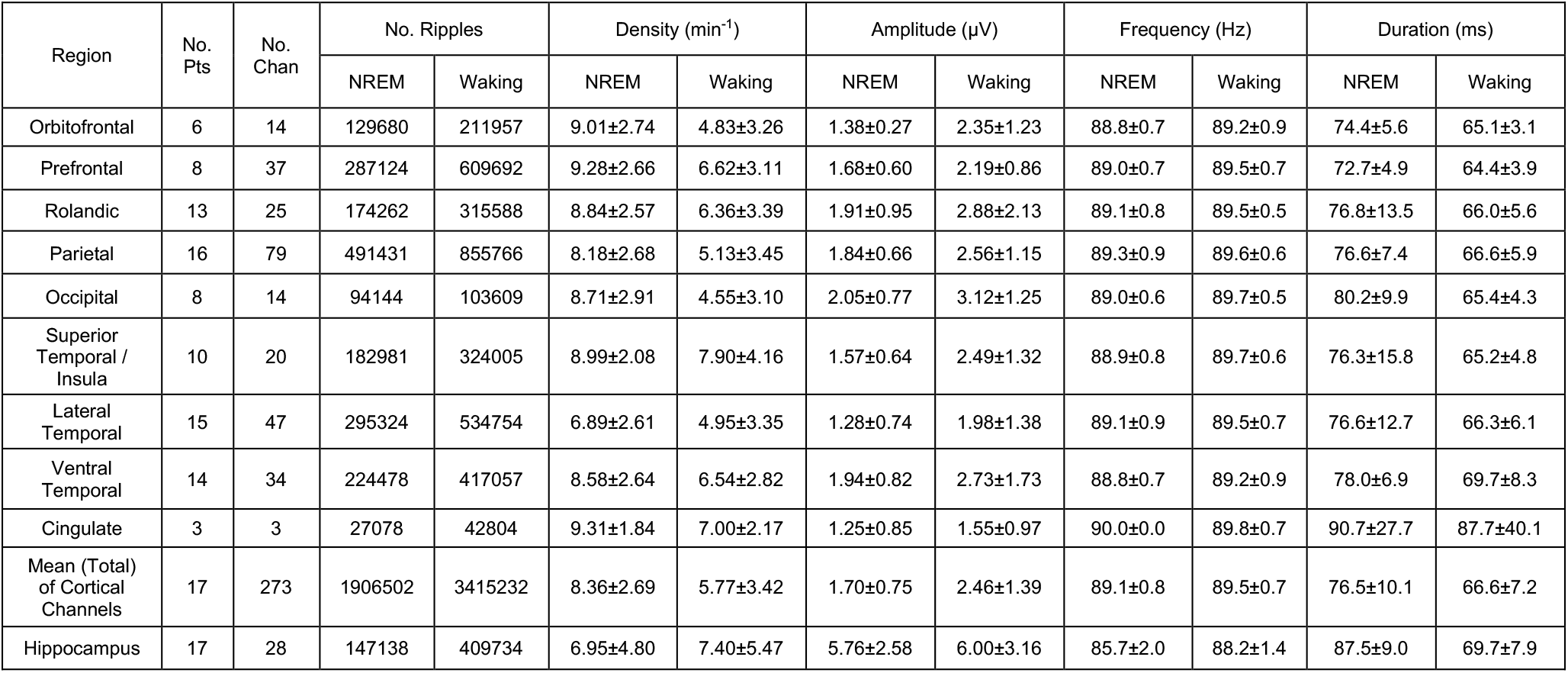
Cortical and hippocampal ripple characteristics by region in NREM and waking. Cortical parcellation scheme is specified in Supplementary Table 2-1. Values are counts or means and standard deviations across channels (*N*=273 cortical, *N*=28 hippocampal) from SEEG patients S1-17.

**Figure 1.**
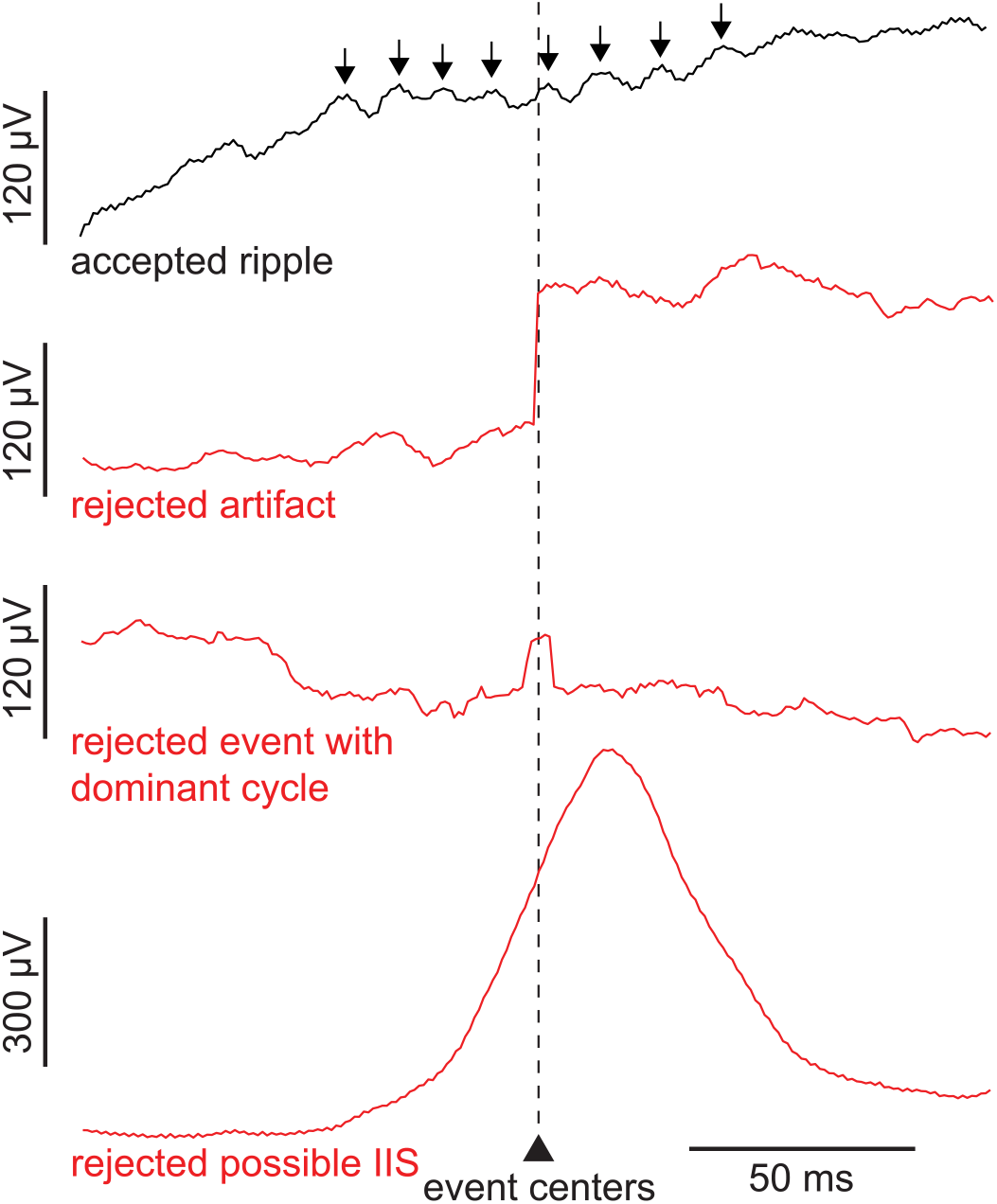
Ripple detection and event rejection. Broadband LFP single sweeps show events that exceeded amplitude thresholds and were accepted for or rejected from the analyses. Traces show an example accepted ripple with black arrows indicating multiple 70-100 Hz oscillation cycles, a rejected artifact, a rejected event with a single dominant cycle, and a rejected possible IIS. Putative ripples within ±500 ms of possible IIS detected on the same channel, as well as putative ripples coinciding with the sharp component of possible IIS on any cortical or hippocampal channel were rejected. IIS=interictal spike, LFP=local field potential.

**Figure 2.**
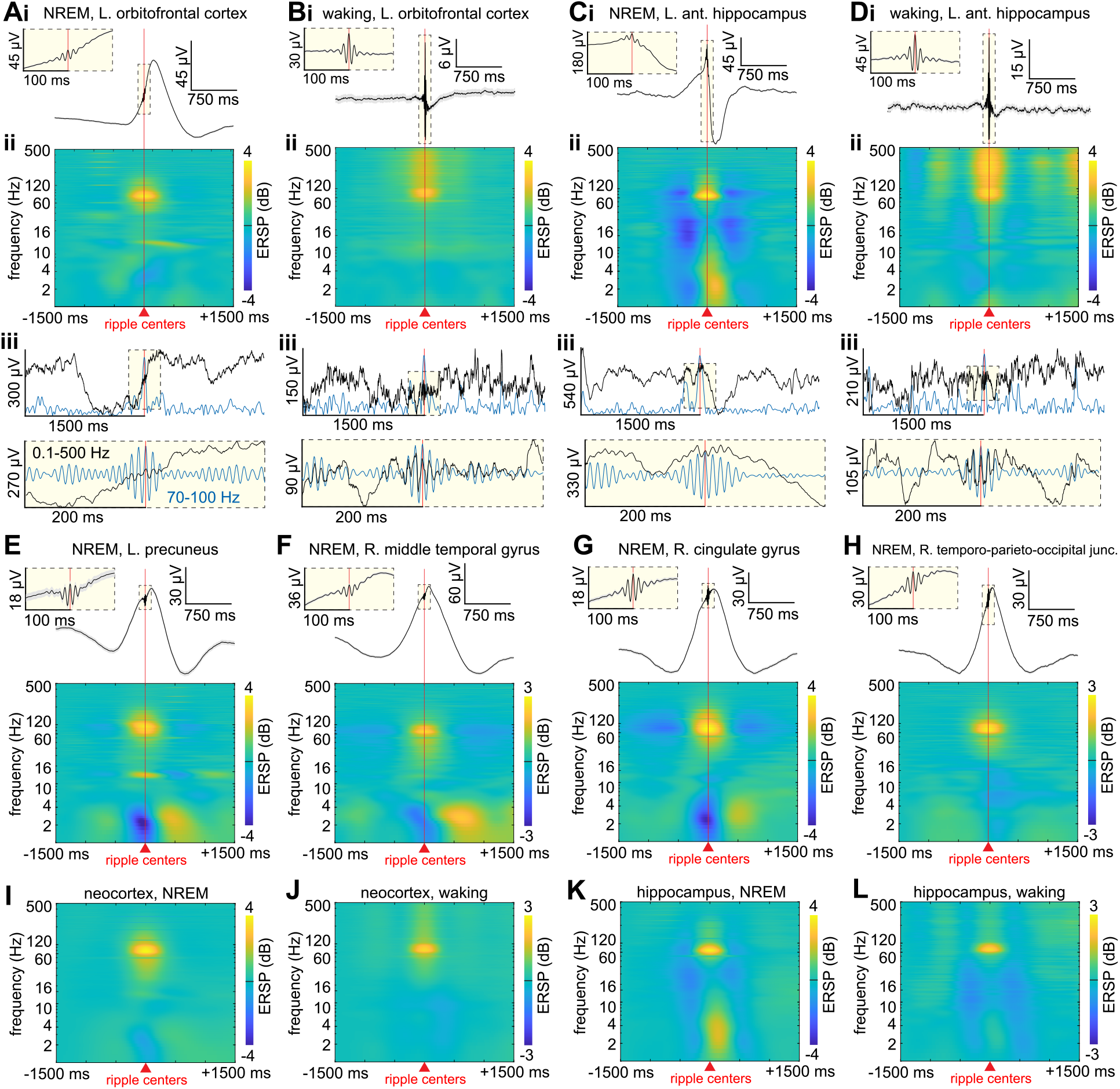
Cortical and hippocampal ripples are generated during NREM and waking. (**A**) Orbitofrontal ripples from one channel during NREM. (**i**) Average broadband LFP. (**ii**) Average time-frequency plot. (**iii**) Example broadband unfiltered 3 s sweep (black) with 70-100 Hz bandpass analytic amplitude (blue) and below the same example, 400 ms sweep (black) with 70-100 Hz bandpass (blue). (**B**) Same as (A) except during waking. (**C**) Same as (A) except hippocampal ripples (note ripple on sharpwave peak). (**D**) Same as (C) except during waking. (**E-H**) Same as (A) except from other cortical regions. (**I-L**) Grand average time-frequency plots across all channels in neocortex (*N*=273) during NREM (**I**) and waking (**J**), as well as in hippocampus (*N*=28) during NREM (**K**) and waking (**L**). Note the highly consistent and focal concentration of power centered at ∼90 Hz, the occurrence of cortical ripples on the down-to-upstate transition during NREM, and that some channels show increased power in the 10-16 Hz spindle band coinciding with the ripples. All plots show ripples detected on bipolar SEEG channels. ERSP=event-related spectral power, NREM=non-rapid eye movement sleep, SEEG=stereoelectroencephalography.

Across states and cortical regions, ripple frequency was remarkably consistent at ∼90 Hz. Specifically, mean cortical ripple frequency within-region ranged from 88.8 to 90.0 Hz during NREM and 89.2 to 89.8 Hz during waking (Table 2, Figure 3). The mean and standard deviation ripple oscillation frequency across all cortical channel means during NREM was 89.1±0.8 Hz and during waking was 89.5±0.7 Hz. These data are also presented as histograms of individual ripple characteristics (amplitude, frequency, duration, associated changes in >200 Hz amplitude) in Supplementary Figure 3-1 as well as individual patients in Supplementary Figure 3-2. The basic characteristics of cortical ripples were very similar in a supplemental analysis which only included cortical channels free of interictal spikes (Figure 4A).

**Figure 3.**
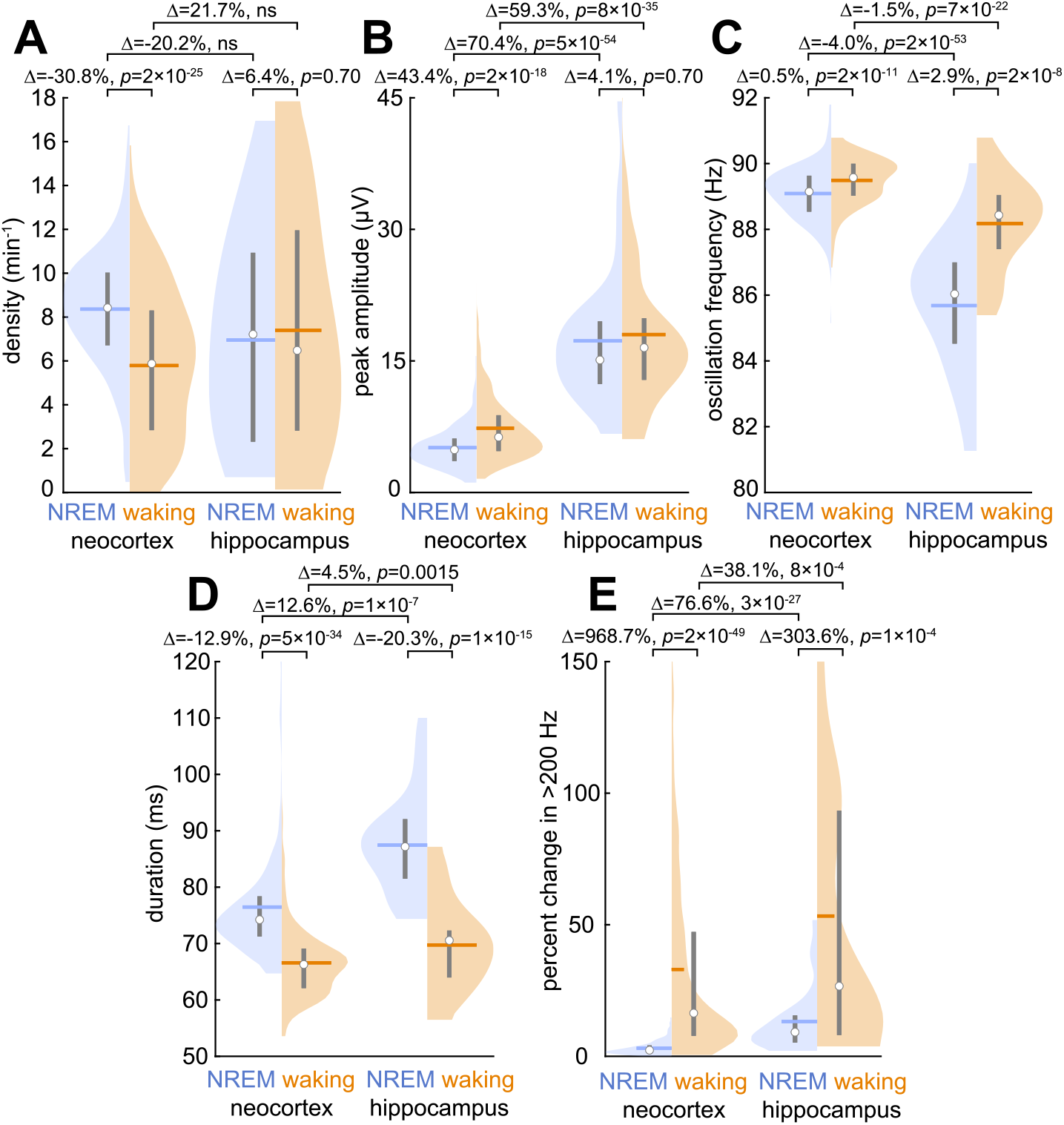
Neocortical and hippocampal ripple characteristics in NREM and waking. (**A-E**) NREM and waking cortical ripple density (**A**), peak 70-100 Hz analytic amplitude (**B**), oscillation frequency (**C**), duration (**D**), and percent change in mean >200 Hz analytic amplitude during ripples compared to a -2 to -1 s baseline (**E**). Distributions are comprised of channel means (SEEG patients S1-17; *N*=273 neocortical channels, *N*=28 hippocampal channels). Circles show medians; horizontal lines, means; vertical lines, interquartile ranges. FDR-corrected *p*-values, linear mixed-effects models with *post hoc* analyses, patient as random effect, ns=non-significant factor precluding *post hoc* analysis. FDR=false discovery rate. See Supplementary Figure 3-1 for distributions across individual ripples and Supplementary Figure 3-2 for distributions across individual patients. Supplementary Figure 3-3 shows that the ∼90 Hz ripple frequency we measured is not due to filtering including the detection bandpass.

**Figure 4.**
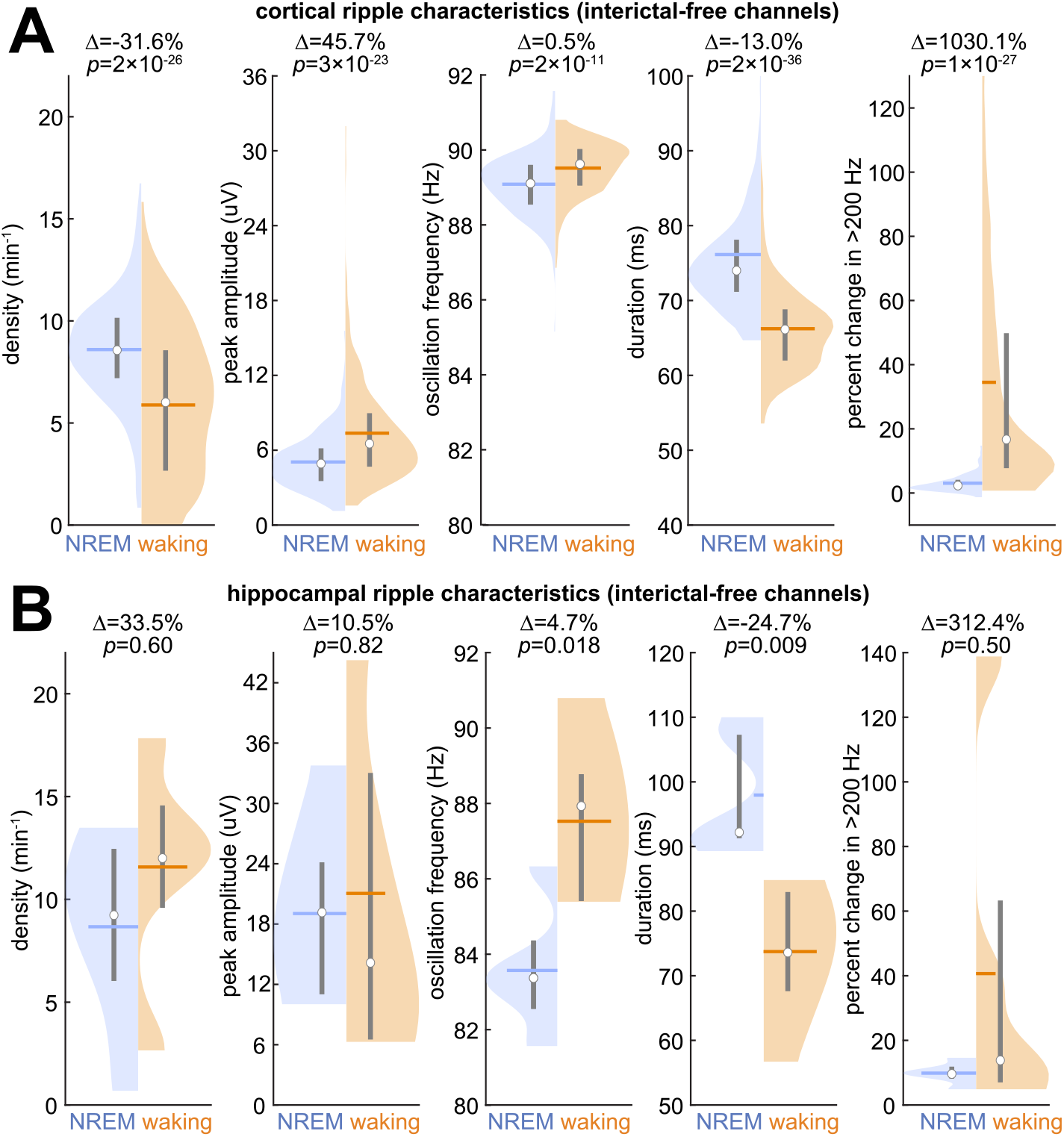
Characteristics of ripples detected on interictal-free channels. (**A**) NREM and waking cortical ripple density, peak 70-100 Hz analytic amplitude, oscillation frequency, duration, and percent change in mean >200 Hz analytic amplitude in interictal-free cortical channels (*N*=232 channels; patients S1-17). (**B**) Same as (A) except interictal-free hippocampal channels (*N*=5 channels; patients S4,6,9,17).

Oscillation frequency was not only highly consistent at ∼90 Hz across ripples, but increased amplitude was highly concentrated during ripples at ∼90 Hz in broadband recordings, with a steep drop-off in amplitude at higher and lower frequencies. This focal power increase, which is especially prominent during NREM, can be seen in example channels (Figure 2A,E-H) as well as grand average time-frequency plots that are averages across all channel averages of all ripples (Figure 2I-L).

We conducted supplementary analyses to examine whether this regularity and focality of ripple frequency could be due to our method of detecting them. First, we conducted a complete re-analysis of all SEEG channels and epochs using two different frequency ranges for ripple detection and selection. One used a detection bandpass from 70-100 Hz as in our primary analysis, and the other expanded the detection bandpass to 65-120 Hz. Mean and standard deviation NREM ripple oscillation frequency across channels (*N*=273) from all SEEG patients (S1-17) is highly similar when using either a 70-100 Hz bandpass (mean±SD=89.1±0.8 Hz) or a 65-120 Hz bandpass (92.7±2.2 Hz) (Supplementary Figure 3-3A). Furthermore, we verified that the 60 and 120 Hz notch filters (to remove line noise) as well as the 70-100 Hz bandpass did not artificially produce a peak in power at ∼90 Hz (Supplementary Figure 3-3B). Thus, the ripple oscillation frequency of ∼90 Hz appears to be physiologic rather than being driven by specific detection methods.

The mean ripple duration was also remarkably consistent (∼70 ms) across states and cortical regions (Table 2). This duration and consistency is not explained by the detection requirement of having at least 3 cycles (which at 90 Hz is 33.3 ms), suggesting that this duration is also a physiological characteristic. Other characteristics including density (NREM: 8.36±2.69 min^-1^, waking: 5.77±3.42 min^-1^), peak amplitude (NREM: 5.10±2.25 µV, waking: 7.38±4.17 µV), and change in >200 Hz amplitude (NREM: 3.09±2.75%, waking: 33.02±36.90%) were also highly similar across channels (Figs.2,3). Ripple charactereristics were also consistent across patients (Supplementary Figure 3-2). However, small but significant differences were noted between regions, cortex vs. hippocampus, and NREM vs. waking, which are described below.

### Ripples exhibit small but significant differences between cortex and hippocampus

Human hippocampal ripples have previously been selected using a variety of methods (reviewed in Jiang et al. (2019c)). In our previous studies we required that ripples be superimposed on sharpwaves (characteristic of anterior hippocampus (Jiang et al., 2019a)) or spindles (characteristic of posterior hippocampus (Jiang et al., 2019b)). In the current study, we re-analyzed these data using exactly the same criteria and procedures as we used for detection and selection of cortical ripples, so as to avoid any methodological ambiguities, i.e., we did not require the presence or absence of any associated lower frequency LFP signature. Using the same detection criteria, we found that cortical and hippocampal ripples share the same basic characteristics with relatively small differences. The basic hippocampal ripple characteristics were very similar in a supplemental analysis which only included hippocampal channels free of interictal spikes (Figure 4B). During NREM, hippocampal ripples had average time-frequency plots (Figure 2C,K) that resembled the corresponding cortical plots (Figure 2A,E-I) in having concentrated oscillatory activity at ∼90 Hz, but differed in being superimposed on the peak of a local sharpwave-ripple rather than occurring just before the local upstate as is seen in the cortex. During waking, average time-frequency plots of hippocampal (Figure 2D,L) and cortical (Figure 2B,J) ripples again show concentrated oscillatory activity at ∼90 Hz, but also with activity that stretches into higher frequencies as noted above. Statistical analyses (Figure 3) revealed several differences which, although small, were nonetheless significant, attributable to the large numbers of channels. Specifically, compared to cortical ripples, hippocampal ripples were on average: slightly less dense during NREM, and more dense during waking; slightly lower frequency, especially during NREM (but still <3 Hz difference); longer duration, especially during NREM (but <10 ms difference); larger amplitude (by ∼2.5x); and accompanied by a smaller but still significant increase in >200 Hz amplitude during waking compared to NREM. In addition, cortical ripple characteristics were not significantly correlated with the hippocampal connectivity density to their local parcel (Figure 5). Overall, cortical and hippocampal ripples appear to be very similar, except notably in their amplitudes and associated slower waves during NREM (Figure 6A-F).

**Figure 5.**
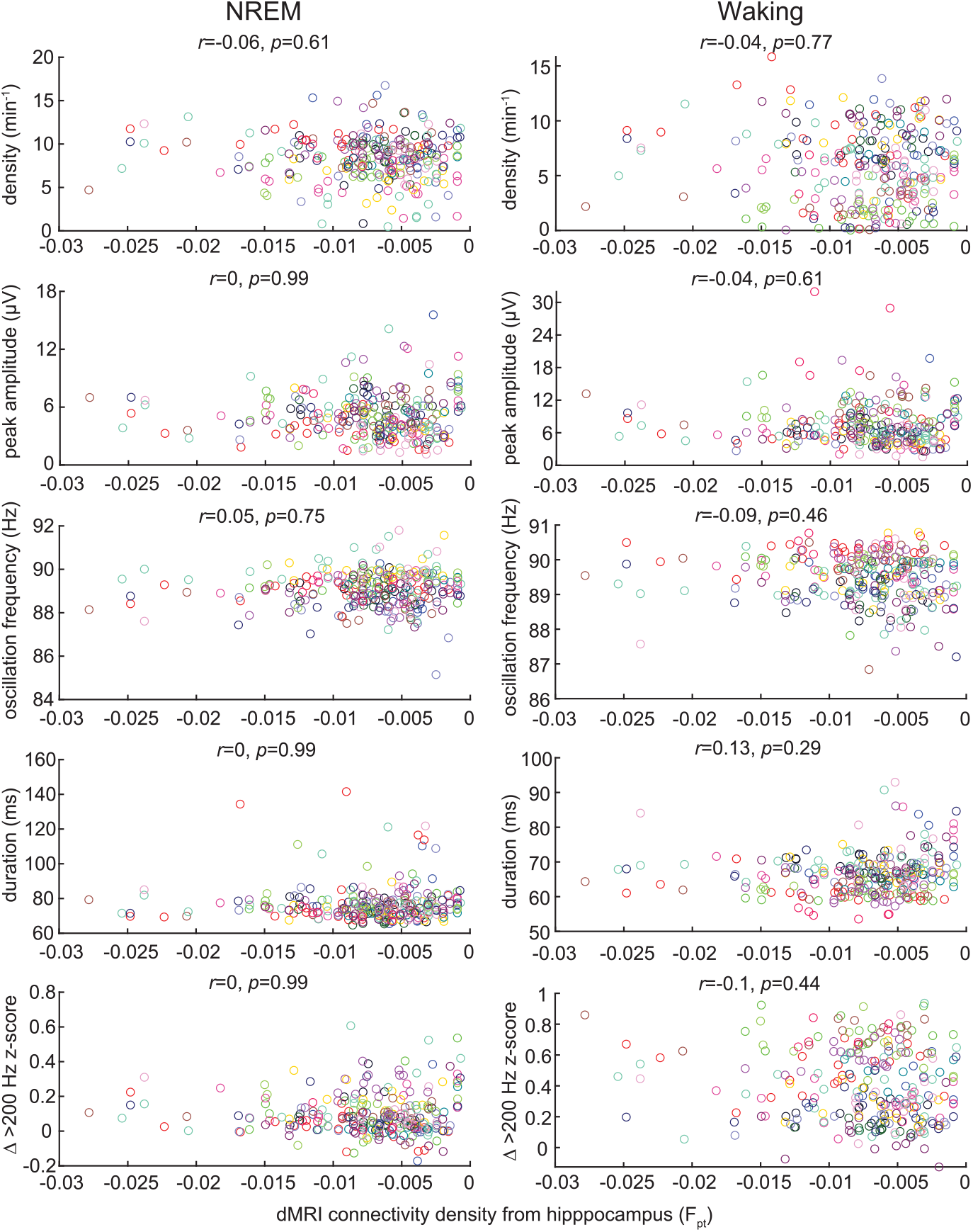
Cortical ripple characteristics versus estimated hippocampo-cortical connectivity density. Cortical ripple density, peak 70-100 Hz analytic amplitude, oscillation frequency, duration, and >200 Hz amplitude modulation (*N*=273 channels from patients S1-17) as a function of the connectivity density (Rosen and Halgren, 2021) between the hippocampus and each cortical parcel. The absence of significant correlations suggests that cortical ripples are related to cortico-cortical integration rather than being driven primarily by the hippocampus.

**Figure 6.**
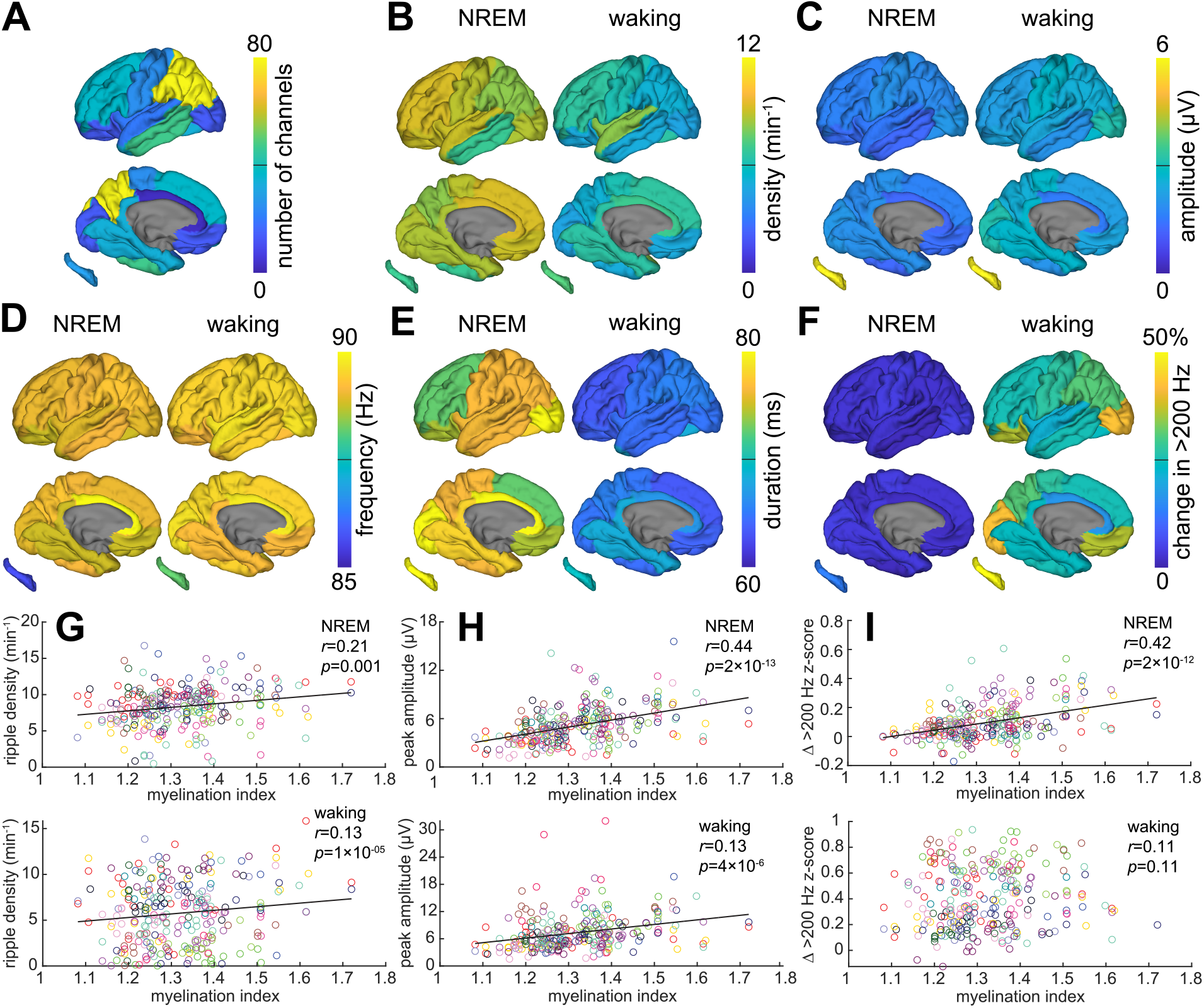
Distributions of ripple characteristics across the cortex and hippocampus in NREM and waking. (**A-F**) Cortical maps with hippocampal map insets of channel coverage (**A**) as well as NREM and waking mean ripple densities (**B**), peak 70-100 Hz analytic amplitudes (**C**), oscillation frequencies (**D**), durations (**E**), and changes in mean >200 Hz analytic amplitude during ripples compared to baseline (−2 to -1 s) (**F**) (SEEG patients S1-17; *N*=273 channels). Left and right hemisphere channels were mapped onto a left hemisphere template. Note that the parcellation scheme (specified in Supplementary Table 2-1) is low resolution and more extensive sampling may detect more spatially-differentiated response characteristics. Values for each cortical region are reported in Table 2. (**G-H**) Average cortical ripple density (**G**) and amplitude (**H**) by channel were significantly correlated with cortical parcel myelination index (i.e., lower densities and amplitudes in association compared to primary areas (Rosen and Halgren, 2021)) during NREM and waking (linear mixed-effects models with patient as random effect, FDR-corrected *p*-values; colors correspond to individual patients). (**I**) Same as (G-H) except change in mean >200 Hz amplitude z-scores, for which there was a significant correlation during NREM but not waking.

### Ripples can be distinguished from high frequency limbic oscillations

Oscillatory activity above 60 Hz in humans has been studied previously, mainly in the hippocampus in relation to epilepsy. Le Van Quyen et al. (2010) reported likely non-pathological activity, mainly from parahippocampal gyrus sites. They imposed a minimum duration of 100 ms, which would have eliminated the vast majority of the events that we studied (Figure 3D; Supplementary Figure 3-1), and is longer than those previously reported for human hippocampal ripples (Jiang et al., 2019c), human cortical ripples (Vaz et al., 2019), and rodent hippocampal ripples (Buzsaki, 2015). Besides being on average ∼7x longer than the events we describe here, those described by Le Van Quyen et al. (2010) also differ in that they contain several oscillatory frequencies (whereas those described here are strongly centered at ∼90 Hz) and have larger amplitudes. Their events are similar, however, in being related to upstates, and in modulating unit firing (see below). We were therefore interested in determining the relationship between the mainly parahippocampal events described by Le Van Quyen et al. (2010) and the widespread cortical events described here. We subselected our parahippocampal channels (*N*=5 channels from 5 patients), and implemented the selection criteria described by Le Van Quyen et al. (2010): data were bandpassed at 40-120 Hz, minimum event duration was 100 ms, and otherwise events were detected as described in the Materials and Methods. Using these selection criteria, we found very low densities of events. The average ripple density of parahippocampal channels during NREM using our criteria was 9.08 min^-1^ (range: 7.23-10.77 min^-1^) whereas that using the Le Van Quyen et al. (2010) detection criteria was only 0.40 min^-1^ (range: 0-1.53 min^-1^). Thus, the events described here do not correspond to those previously reported by Le Van Quyen et al. (2010).

### Ripples exhibit small but significant differences between primary sensory-motor and association cortices

Previous work in sleeping rats (Khodagholy et al., 2017) and waking humans (Vaz et al., 2019) found cortical ripples to be more common in association areas than in early sensory and motor regions where they were absent or infrequent. In contrast, we found that ripple density, amplitude, and accompanying >200 Hz amplitude (a proxy for unit-firing (Mukamel et al., 2005)) in primary cortex were all significantly higher than in association areas, as indicated by a positive correlation with the myelination index (Figure 6G-I) (Rosen and Halgren, 2021). Oscillation frequency and duration were not correlated with myelination index (Figure 7). In some cases, this effect was small, e.g., explaining only 1.7% of the variance in density during waking and 4.4% during NREM. In other cases, the effect was substantial, e.g., explaining 19.4% of the variance in amplitude during NREM. Comparing NREM and waking, in NREM there was a significantly stronger correlation between myelination index and density (*p*=0.02, *t*=2.1), amplitude (*p*=8×10^−4^, *t*=3.2), and >200 Hz modulation (*p*=2×10^−6^, *t*=4.7; one-sided paired *r*-test; MATLAB: *r_test_paired* (Steiger, 1980)). In sum, human cortical ripples appear to be modestly more numerous and robust in sensory and motor than association areas.

**Figure 7.**
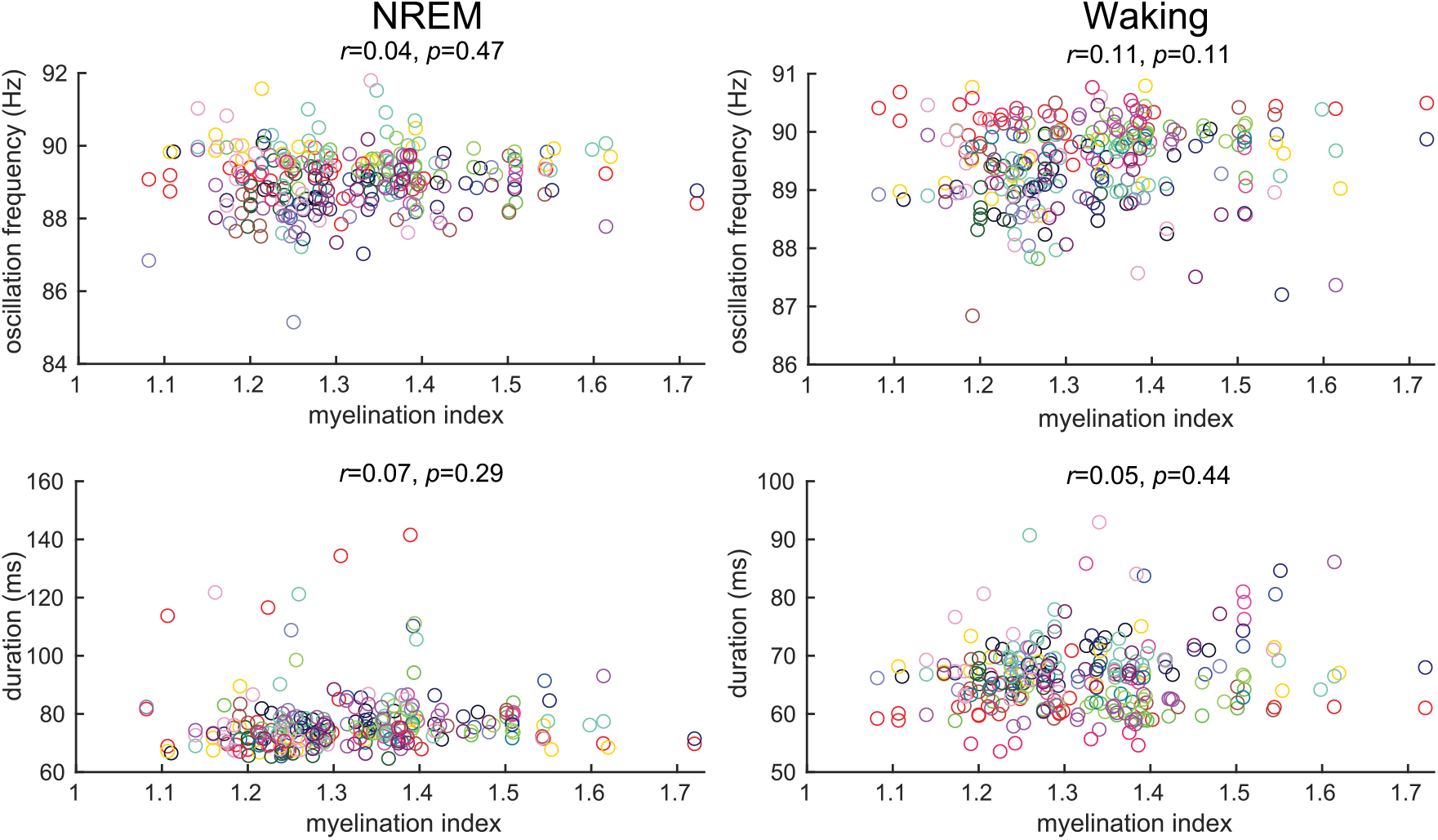
Cortical ripple oscillation frequency and duration versus myelination index. Neither cortical ripple oscillation frequency nor duration were significantly correlated with myelination index (Rosen and Halgren, 2021) during NREM or waking (linear mixed-effects models with patient as random effect, FDR-corrected *p*-values, *N*=273 channels from patients S1-17). Colors correspond to individual patients. Of note, higher myelination indices correspond to primary whereas lower myelination indices correspond to associative cortical regions.

### Ripples exhibit small but significant differences between NREM and waking

Most cortical and hippocampal ripple characteristics were similar between NREM and waking (Figure 3, Table 2), but due to the large numbers of events, even small differences were significant. Considering only differences >10%, cortical ripple density was 31% higher and amplitude was 43% lower during NREM than waking. Ripple duration was 13% lower in the cortex and 20% lower in the hippocampus during NREM than waking. The most striking differences were the percent change in mean >200 Hz amplitude during ripples compared to a -2 to -1 s baseline (Figure 3E), which increased 969% from NREM to waking in the cortex, and 304% in the hippocampus. This difference is clearly seen to be broadband in the waking ripple triggered time-frequency plots from the cortex (Figure 2Bii,J) and hippocampus (Figure 2Dii,L), and thus would typically be interpreted not as an oscillation, but as an indication of increased multi-unit activity. Thus, compared to waking, cortical ripples are somewhat denser, longer, and smaller in NREM; hippocampal ripples are also shorter. The only major difference is that both cortical and hippocampal ripples appear to be associated with a much larger increase in putative multi-unit activity during waking than NREM.

### Cortical ripples lock to sleep waves crucial for memory consolidation

Oscillation couplings are important for memory consolidation (Latchoumane et al., 2017). Previous studies have shown that cortical ripples occur on upstate and spindle peaks in cats (Grenier et al., 2001) and rats (Khodagholy et al., 2017), but these relationships have not been evaluated in humans. We detected downstates and upstates, with polarities determined based on associated high frequency activity (see the Materials and Methods for details), as well as spindles, and found that cortical ripples were precisely coupled to the sequence of sleep waves described above (Figure 8). Specifically, in 95% of cortical channels, ripples were significantly associated with downstates and upstates (Figure 8B,D,E), usually on the down-to-upstate transition, as seen in individual trials (Figure 2Aiii; Figure 8A), and in ripple-triggered averages of the broadband LFP (Figure 2Ai,E-H). Peri-ripple histograms show that, on average, cortical ripple centers occurred 450 ms after downstate maxima and 100 ms before upstate maxima (Figure 8B,D-G). Less frequently, ripples occurred during spindles (significant association in 29% of channels; Figure 8C,E), as seen in individual trials (Figure 8A), and in peri-ripple time-frequency plots (Figure 2Aii,E). Peri-ripple histograms show that spindles began on average 225 ms prior to the cortical ripple center, indicating that ripples tend to occur during spindles (Figure 8C,E-G). The probability of ripples occurring during spindles preceding upstates was greater than that for ripples occurring during spindles, or before upstates (Figure 8H). These results suggest that the timing of cortical ripples during NREM is appropriate for facilitating consolidation, guided by a sequential activation of sleep waves.

**Figure 8.**
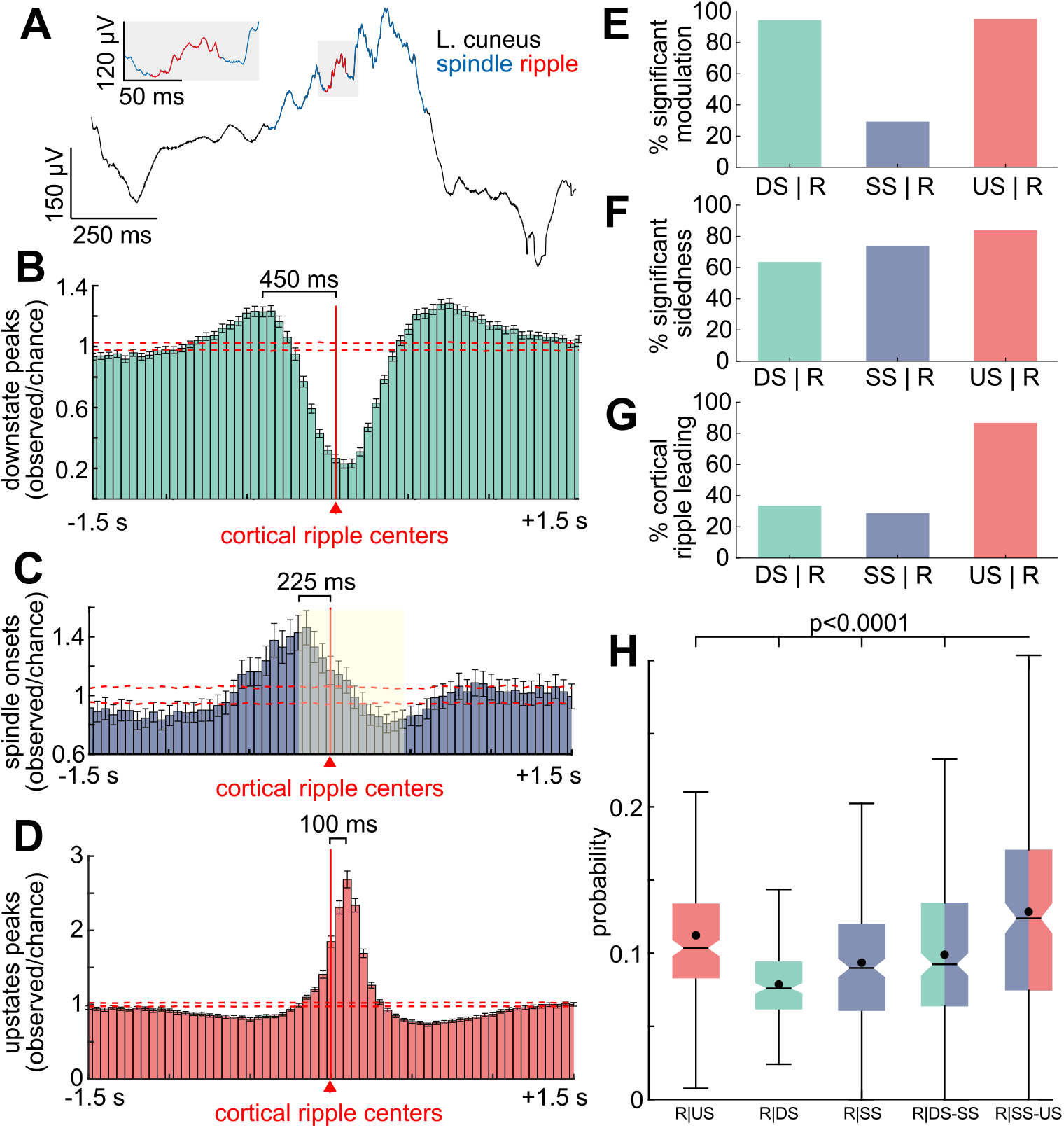
Cortical ripples occur during sleep spindles on the down-to-upstate transition. (**A**) Example cortical ripple occuring during a spindle on a down-to-upstate transition during NREM. (**B**) Times of downstate peaks plotted relative to local cortical ripple centers at *t*=0 across significant channels (*N*=258/273, patients S1-17) during NREM. Downstate maxima occurred on average 450 ms prior to the cortical ripple center. Dashed red lines show 99% confidence interval of the null distribution (200 shuffles/channel). (**C**) Same as (B) except spindle onsets (*N*=80/273). Spindles began on average 225 ms prior to the cortical ripple center, indicating that ripples occurred during spindles (yellow shaded area indicates average spindle interval of 634 ms). (**D**) Same as (B) except upstate peaks (*N*=260/273). Upstate maxima occurred on average 100 ms after the cortical ripple center. (**E**) Percent of channels with significant peri-ripple modulations of sleep waves detected on the same channels within ±1000 ms (e.g., US | R represents upstate peaks relative to cortical ripples at *t*=0; one-sided randomization test, 200 shuffles, 50 ms non-overlapping bins, 2 consecutive bins with post-FDR *p*<0.05 required for significance). DS and US were significantly associated with ripples in ∼95% of channels, sleep spindles less frequently. (**F**) Percent of channels with significant modulations that had significant sidedness preference around *t*=0 (post-FDR *p*<0.05, one-sided binomial test, -1000 to -1 ms vs. 1 to 1000 ms, expected=0.5). (**G**) Percent of channels with significant sidedness around 0 that had cortical ripples leading the other sleep waves (according to counts in -1000 to -1 ms vs. 1 to 1000 ms). Downstate peaks and spindle onsets typically preceded ripples, and upstate peaks followed. (**H**) Probabilities of ripple centers preceding upstates, following downstates, occurring during spindles in isolation or during spindles following downstates (DS-SS), or during spindles preceding upstates (SS-US). The time window used following a downstate or preceding an upstate was 634 ms, which was the average spindle duration. Note the probability of a ripple occurring was greatest during spindles preceding upstates (post-FDR *p<*0.0001, two-sided paired *t*-test, channel-wise). See Supplementary Figure 8-1 for results from (E-G) in tabular format. DS=downstate, SS=sleep spindle, US=upstate.

### Pyramidal cell firing precedes interneuron firing at cortical ripple peaks

Using microelectrode array recordings from granular/supragranular layers of lateral temporal cortex during NREM, we detected ripples (Figure 9A) as well as action-potentials, which were sorted into those arising from putative single pyramidal or interneuron units. The mean and standard deviation ripple oscillation frequency across microelectrode channels (*N*=72) was 89.6±1.4 Hz, and across individual ripples (N=50,967) was 90.1±5.2 Hz. We found that interneurons had a strong tendency to fire at the peak of the ripple, whereas pyramidal cells fired shortly before (Figure 9B). We have previously shown that, in humans, upstates (Csercsa et al., 2010) and spindles (Dickey et al., 2021) are associated with increased unit-firing rates. Since cortical ripples are precisely coupled to these events (Figure 8), their occurrence on upstates and spindles implies an underlying phasic depolarization, which can generate ∼90 Hz oscillations in computational and experimental models via pyramidal-interneuron feedback inhibition (Bazhenov et al., 2008; Buzsaki, 2015). Depolarization causes basket cells to fire synchronously, inhibiting other basket cells and pyramids via GABA_A_. Pyramids fire upon recovery, exciting basket cells as they recover, leading to another cycle. As this model would predict, we found that pyramids and interneurons were strongly phase-locked to cortical ripples (Figure 9B), with pyramidal significantly leading interneuron spiking (Figure 9C-D). Furthermore, interneurons fired at the ripple peak, when pyramidal somatic inhibition would be maximal, as found in cats (Grenier et al., 2001). Similarly, ripple amplitude was higher in waking than NREM (Figure 3B; Figure 6C), consistent with relative depolarization of pyramidal membrane potential during waking. Phasic depolarization during waking ripples was also suggested by increased >200 Hz amplitude (Figure 2B,J; Figure 3E; Figure 6F). Thus, human cortical ripples are associated with strong tonic and phase-locked increases in pyramidal and interneuron firing, and are likely generated by a sudden depolarization triggering pyramidal-interneuron feedback oscillations (Figure 9E).

**Figure 9.**
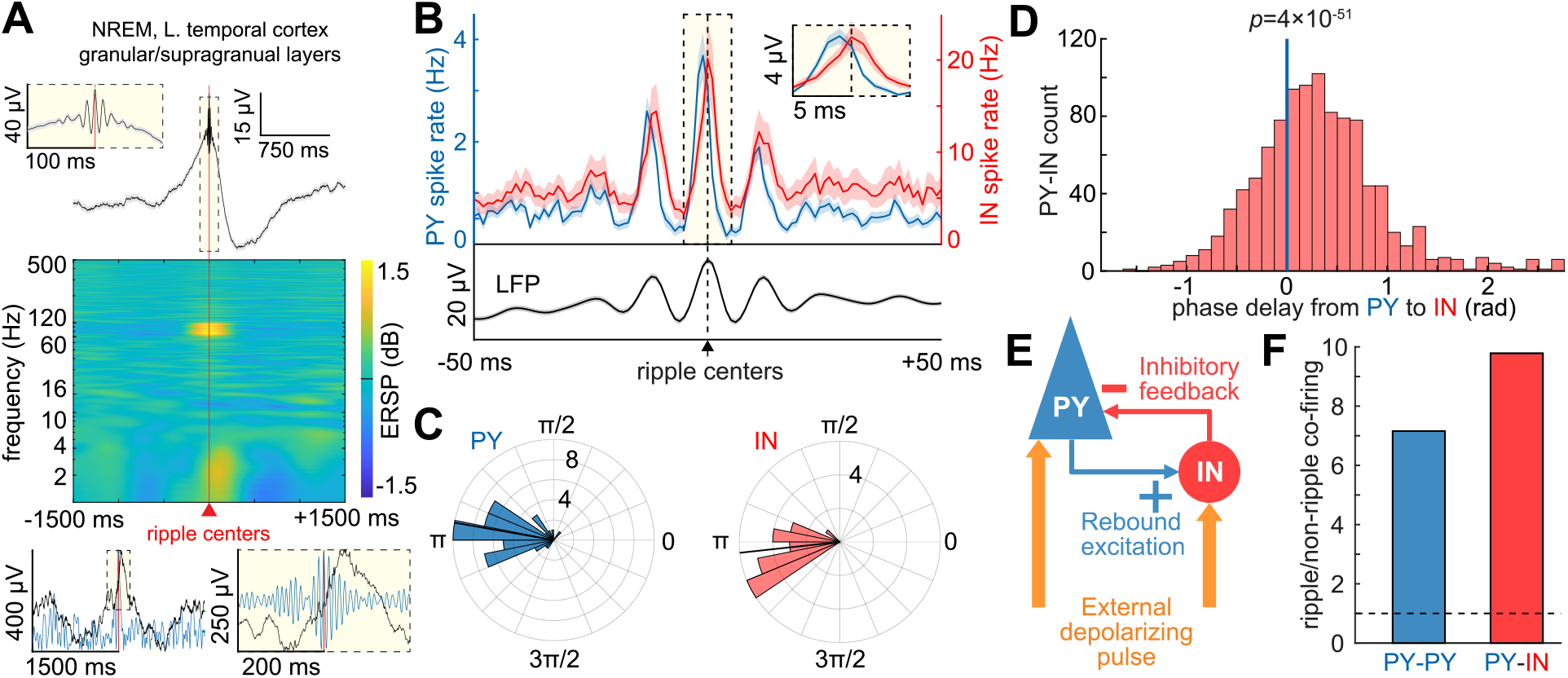
Pyramidal leads interneuronal cell firing at cortical ripple peaks. (**A**) Superior temporal gyrus granular/supragranular layer ripples detected during NREM in a Utah Array recording. Top shows average broadband LFP, middle shows average time-frequency, and bottom shows single trial example trace in broadband LFP (black) and 70-100 Hz bandpass (blue, left–analytic amplitude, right–sweep). (**B**) Mean broadband LFP locked to cortical ripple centers (black) and associated local PY (blue; *N*=69) and IN (red; *N*=23) spike rates during NREM. (**C**) Circular mean 70-100 Hz phase of spikes of each PY (*N*=47, mean=2.97 rad, *p*=5×10^−21^, Rayleigh test) and IN (*N*=22, mean=3.25 rad, p=3×10^−13^) during local cortical ripples (minimum 30 spikes per unit). PY spiking preceded IN spiking by 0.28 rad (*p*=0.02, Watson-Williams test). Note that 0 rad corresponds to the trough and π rad corresponds to the peak of the ripple. (**D**) Circular mean ripple phase-lags of spikes from each PY (*N*=47) to each IN (*N*=22) (*N*=1034 unit pairs, mean=0.31±0.63 rad, *p*=4×10^−51^, one-sided one-sample *t*-test). (**E**) Pyramidal Interneuron Network Gamma ripple generation mechanism consistent with single-unit recordings, animal studies (Stark et al., 2014), and modelling (Buzsáki and Wang, 2012). Abrupt depolarization causes synchronous PY and IN firing, which then spike rhythmically separated by fixed intervals due to recurrent inhibition. (**F**) Pairs of pyramidal cells (PY-PY) co-fire ∼7 times more often within 5 ms of each other during ripples compared to randomly selected epochs in between ripples matched in number and duration to the ripples. Similarly, pyramidal-interneuron (PY-IN) pairs co-fire ∼10 times more during ripples. IN=putative interneuron unit, PY=putative pyramidal unit.

### Cortical ripples group co-firing with timing optimal for spike-timing-dependent plasticity

The increased tonic and phase-locked unit firing and occurrence of ripples on spindles on the down-to-upstate transition suggests that ripples may facilitate cortical plasticity (Dickey et al., 2021). This could occur through short latency co-firing between units that leads to STDP. Indeed, we found a large increase during ripples of short latency (<5 ms) co-firing between pyramidal-pyramidal (PY-PY) unit pairs and pyramidal-interneuron (PY-IN) unit pairs (Figure 9F). Specifically, there was an increase in co-firing during ripples vs. non-ripples (PY-PY: 7.16 fold increase, *p*=2×10^−159^, *N*=4638, *t*=28.0; PY-IN: 9.78 fold increase, *p*=8×10^−99^, *N*=1561, *t*=22.7; one-sided paired *t*-test). Non-ripple comparison periods were randomly selected epochs between ripples matched in number and duration.

In order to test if this increase in co-firing was not simply due to an increase in the overall firing rates of units during ripples, or even their rhythmicity, but rather a specific organization of co-firing by the phase of individual ripple cycles, we constructed a control dataset consisting of the neuronal firing during ripples where the ripple times were randomly shuffled. Compared to this shuffled control, there was a significant increase in short-latency co-firing during ripples with their actual spike times (PY-PY: 1.21 fold increase, *p*=1×10^−7^, *N*=4638, *t*=5.2; PY-IN: 1.15 fold increase, *p*=1×10^−7^, *N*=1561, *t*=5.2, one-sided paired *t*-test). This finding indicates that the increase in co-firing is due to a specific organization of co-firing by the phase of the individual ripple cycles. Thus, ripples create a necessary and sufficient condition for STDP, and therefore may underlie the crucial contribution of these nested waves to consolidation (Dickey et al., 2021).

## Discussion

The current study provides the first comprehensive characterization of cortical ripples during waking and NREM in humans. Using intracranial recordings, we report that human cortical ripples are widespread during NREM in addition to waking. Basic ripple characteristics (occurrence rate, amplitude, oscillation frequency, duration, distribution across cortical areas) are stereotyped across electrodes and cortical parcels. They are also very similar in cortex vs. hippocampus, and in NREM vs. waking. Cortical ripple stereotypy and ubiquity across structures and states suggests that they may constitute a distinct and functionally important neurophysiological entity.

The characteristics of human ripples are similar to those of rodents with two exceptions. First, the center frequency of human ripples is focally ∼90 Hz (Jiang et al., 2019a), whereas in rodents it is ∼130 Hz (Khodagholy et al., 2017). Conceivably, the larger human brain may require more time for cortico-cortical and hippocampo-cortical integration than the rodent brain. The second apparent difference between humans and rodents is in cortical distribution. A previous rodent study found that cortical ripples were restricted to association cortex (Khodagholy et al., 2017), and consistent observations were reported in humans (Vaz et al., 2019). In contrast, we recorded cortical ripples in all areas sampled, with slightly lower occurrence rates in association areas as indicated by the positive correlation between occurrence rate and myelination index.

In contrast to the similarities of basic properties, we found that ripples in waking vs. NREM occur within much different immediate physiological contexts. In NREM, cortical ripples are strongly associated with local downstates and upstates, and less strongly with sleep spindles. Characteristically, ripples occur on the upslope ∼100 ms before the peak of the upstate. Previous studies found that sleep spindles (Dickey et al., 2021) and upstates (Csercsa et al., 2010) are associated with strong increases in local unit-firing, and we found that also to be the case for cortical ripples during NREM. Hippocampal ripples during NREM in previous studies were also found often to occur during local sharpwaves or spindles (Staresina et al., 2015; Jiang et al., 2019a; Jiang et al., 2019b; Jiang et al., 2019c).

Since sleep spindles and down-to-upstates are characteristic of NREM, they would not be expected to occur with waking ripples. Although we found no other lower-frequency wave to be consistently associated with ripples during waking, both cortical and hippocampal ripples occurred during greatly increased local >200 Hz amplitude activity, a surrogate for unit-firing (Mukamel et al., 2005). Thus, the local contexts of both cortical and hippocampal ripples, in both NREM and waking, are sudden increases in local excitability lasting at least as long as the ripple (∼70 ms). NREM and waking are different in that the depolarizing pulse is organized by endogenous sleep rhythms in NREM, but appears to be related to exogenous input, such as a retrieval cue, during waking.

Computational neural models and experimental studies have found that depolarizing pulses can induce oscillations from ∼20-160 Hz (Buzsáki and Wang, 2012; Buzsaki, 2015). Depolarization causes synchronous pyramidal cell firing, which is inhibited for a fixed time by recurrent inhibition from local basket cells. The pyramidal cells then fire again, resulting in Pyramidal Interneuron Network Gamma (PING). As predicted by this model, we found strong ripple phase-modulation of putative pyramidal and interneuronal firing, with pyramids regularly preceding interneurons. Our findings are also consistent with PING supplemented by synchronized basket cell firing and mutual inhibition (ING; Interneuron Network Gamma) (Bartos et al., 2002). Definitive mechanistic demonstration would require experiments not currently possible *in vivo* in humans (Stark et al., 2014).

Previous studies have shown that cortical ripples occur on upstate and spindle peaks in cats (Grenier et al., 2001) and rats (Khodagholy et al., 2017), and parahippocampal gamma bursts of various frequencies occur on upstates in humans (Le Van Quyen et al., 2010). Here we report similar relationships in humans, consistent with a previous finding that cortical ripples during human NREM are suppressed during and increased following downstates (von Ellenrieder et al., 2016). The critical role of rodent hippocampal ripples in memory consolidation during NREM (Girardeau et al., 2009) is dependent on their association with cortical sleep waves (spindles, downstates, upstates) (Siapas and Wilson, 1998; Maingret et al., 2016; Latchoumane et al., 2017). In humans hippocampal ripples are also associated with sleep waves (Staresina et al., 2015; Latchoumane et al., 2017; Jiang et al., 2019a; Jiang et al., 2019b), and cortical sleep waves are also associated with consolidation (Niknazar et al., 2015). Thus, our observation that human cortical ripples during NREM are strongly associated with upstates, and less strongly with downstates and spindles, is consistent with a role of human cortical ripples in consolidation. This is supported by a recent study showing that cortical ripples during NREM mark the replay of learned motor patterns from prior waking (Rubin et al., 2022).

Consolidation requires plasticity to increase the strength of the connections embodying the memory, which may occur when pre- and post-synaptic cells fire in close temporal proximity, termed STDP (Feldman, 2012). We show here that local neurons are much more likely to fire at delays optimal for STDP during ripples than control periods, and that co-firing is organized beyond what would be expected by a general increase in neuron firing. A similar increase is also observed during sleep spindles and upstates in humans (Dickey et al., 2021). Since we also show that NREM cortical ripples are temporally coordinated with sleep spindles and upstates, this supports a synergistic facilitation of plasticity. Thus, multiple characteristics of cortical ripples are consistent with them playing a role in consolidation, but direct confirmation of this role will require interventions that were not performed in the current study.

Recently, human waking cortical ripples were shown to mark spatiotemporal firing patterns during cued recall of items that reproduced those previously evoked by the same items during their initial presentation (Jiang et al., 2017). This suggests the possibility that ripples, in humans and rodents, NREM and waking, hippocampus and cortex, share a common role in contributing to the reconstruction of previously occurring firing patterns. The similar characteristics of human ripples, in NREM and waking, hippocampus and cortex, is consistent with this speculation.

Previous reports in rodents (Khodagholy et al., 2017) and humans (Vaz et al., 2019) that cortical ripples were restricted to association cortex were also intertpreted as consistent with a selective interaction of cortical ripples with the hippocampus. However, in our more extensive sample of cortical sites, we observed a lower ripple density in higher associative cortex. Furthermore, we found no relationship across cortical parcels between their degree of connectivity with the hippocampal formation (inferred from diffusion imaging (Rosen and Halgren, 2021)) and cortical ripple density or any other ripple characteristic. Indeed, in other work, we found that cortical ripples are more likely to co-occur and phase-lock with other cortical sites than the hippocampus (Dickey et al., 2022). Thus, the functional role of cortical ripples may not be confined to memory consolidation and recall.

In summary, the current study provides the first report of cortical ripples during sleep in humans. Cortical ripples during NREM were found to have basic characteristics (duration, oscillation frequency, and occurrence rate) that are highly similar to those of cortical ripples during waking as well as hippocampal ripples. Cortical ripple oscillation frequencies were tightly focused at ∼90 Hz. This finding, together with their stereotypy and ubiquity across structures and states, suggests that cortical ripples may constitute a distinct and functionally important neurophysiological entity. Unit firing during human cortical NREM ripples supported the mechanism proposed for rodent hippocampal ripples, based on pyramidal-interneuron feedback. Cortical ripples during NREM characteristically occurred after downstates, during spindles, shortly before the peak of the upstate, an association and timing that is important for the consolidation of memories during sleep. Furthermore, NREM cortical ripples were associated with increased local co-firing between units, thus fulfilling a fundamental requirement for STDP. However, cortical ripples were widespread across cortical parcels, regardless of their probable connectivity with the hippocampus. Overall, these characteristics of cortical ripples during human NREM are consistent with a role that includes but may not be limited to sleep-dependent memory consolidation.

## Materials and Methods

### Patient selection

Data from a total of 18 patients (12 female, 30.0±12.2 years old) with pharmaco-resistant epilepsy undergoing intracranial recording for seizure onset localization preceding surgical treatment were included in this study (Table 1). Patients whose SEEG recordings were analyzed were only included in the study if they had no prior brain surgery; background EEG (with the exception of epileptiform transients) in the normal range; and electrodes implanted in what was eventually found to be non-lesional, non-epileptogenic cortex, as well as non-lesional, non-epileptogenic hippocampus (such areas were suspected to be part of the focus prior to implantation, or were necessary to pass through to reach suspected epileptogenic areas).

Furthermore, 1 of these patients (51 year old right handed female) was also implanted with an intracranial microelectrode (Utah Array) into tissue that was suspected based on pre-surgical evaluation to be included within the region of the therapeutic resection. The implantation of the array did not affect clinical monitoring. It was later resected in order to gain access to the surgical focus beneath and the electrode was determined not to be implanted in an epileptogenic zone and no seizures originated from the region of the array.

Patients were excluded from the study if they had prior brain surgery or did not have non-lesioned hippocampal and cortical channels that were not involved in the early stage of the seizure discharge and did not have frequent interictal activity or abnormal LFPs. Utah Array patients were only included in the study if they had at least 20 PY and 20 IN. Based on these criteria, 18 patients were included in this study out of 84. All patients gave fully informed written consent for their data to be used for research as monitored by the local Institutional Review Boards at Cleveland Clinic and Partners HealthCare (including Massachusetts General Hospital).

### Intracranial recordings

Patients were implanted with intracranial electrodes for ∼7 days with continuous recordings for seizure onset localization. SEEG electrode implantation and targeting were made for purely clinical purposes. SEEG recordings were collected with a Nihon Kohden JE-120 amplifier at 1000 Hz sampling (patients S1-17). Standard clinical electrodes were 0.8mm diameter, with 10-16 contacts of length 2mm at 3.5-5mm pitch (∼150 contacts/patient).

Microelectrode recordings from 1 patient implanted with a Utah Array were also analyzed (200 min of NREM). The Utah Array is a 10×10 microelectrode array with corners omitted and 400 µm contact pitch (Waziri et al., 2009; Keller et al., 2010; Fernández et al., 2014). Each silicon probe is 1mm long with a base of 35-75 µm that tapers to 3-5 µm. The probes are insulated except for the platinum-coated tip. Data were acquired at 30 kHz (Blackrock Microsystems) with a 0.3-7.5 kHz bandpass. Data were recorded with respect to a distant reference wire.

### Electrophysiology pre-processing

Offline data preprocessing was performed in MATLAB 2019b and LFPs were inspected visually using the FieldTrip toolbox (Oostenveld et al., 2011). SEEG data were downsampled to 1000 Hz with anti-aliasing and 60 Hz notch filtered (zero-phase) with 60 Hz harmonics up to 480 Hz. Transcortical contact pairs were identified using both anatomical location (using the pre-operative MRI aligned to the post-operative CT), and physiological properties (high amplitude, coherence and inversion of spontaneous activity between contacts), and selected such that no 2 pairs shared a contact. All SEEG analyses were performed using bipolar derivations between adjacent contacts in cortical or hippocampal gray matter in order to ensure that activity was locally generated (Mak-McCully et al., 2015).

### Channel selection

Channels were excluded from analysis if they were in lesioned tissue, involved in the early stages of the seizure discharge, had frequent interictal activity, or abnormal LFPs. From the total 2129 bipolar channels (1202 left hemisphere) of the 17 SEEG patients (S1-17), 28 hippocampal (16 left hemisphere) and 273 transcortical (133 left hemisphere) bipolar channels were selected for the analyses (Table 1). Note that most channels were rejected because they did not constitute a transcortical pair as described above. First, most bipolar pairs were in the white matter, and thus did not record focal cortical activity. In addition, many channels were rejected for the related criterion that one of the contacts was in common with another bipolar pair that was already selected. This was done because a common contact means that the two bipolar pairs would not provide independent measurements. Polarity was corrected for individual bipolar channels such that downstates were negative and upstates were positive. This was accomplished by ensuring that negative peaks during NREM were associated with decreased and positive peaks were associated with increased mean ±100 ms 70-190 Hz analytic amplitude, an index of cell-firing that is strongly modulated by downstates and upstates (Csercsa et al., 2010).

### Electrode localization

Cortical surfaces were reconstructed from the pre-operative whole-head T1-weighted structural MR volume using the standard FreeSurfer recon-all pipeline (Fischl, 2012). Atlas-based automated parcellation (Fischl et al., 2004) was used to assign anatomical labels to regions of the cortical surface in the Destrieux atlas (Destrieux et al., 2010). In addition, automated segmentation was used to assign anatomical labels to each voxel of the MR volume, including identifying voxels containing hippocampal subfields (Iglesias et al., 2015). In order to localize the SEEG contacts, the post-implant CT volume was registered to the MR volume, in standardized 1mm isotropic FreeSurfer space, using the general registration module (Johnson et al., 2007) in 3D Slicer (Fedorov et al., 2012). The position of each SEEG contact, in FreeSurfer coordinates, was then determined by manually annotating the centroids of the electrode contact visualized in the co-registered CT volume. Each transcortical contact pair was assigned an anatomical parcel from the atlas above by ascertaining the parcel identities of the surface vertex closest to the contact pair midpoint. Subcortical contacts were assigned an anatomical label corresponding to the plurality of voxel segmentation labels within a 2-voxel radius. Transcortical contact pair locations were registered to the fsaverage template brain for visualization by spherical morphing (Fischl et al., 1999). To plot values on a template brain, channel means were averaged for each cortical region (with the two hemispheres combined) and then morphed onto a left hemisphere ico5 fsaverage template. White-matter streamline distances between channels were computed using the 360 parcels of the HCP-MMP1.0 atlas (Glasser et al., 2016), as determined by probabilistic diffusion MRI tractography (Behrens et al., 2007), are population averages from Rosen and Halgren (2021). When two channels were in the same HCP parcel, the distance was considered to be 0.

### Time-frequency analyses

Average time-frequency plots of the ripple event-related spectral power (ERSP) were generated from the broadband LFP using EEGLAB (Delorme and Makeig, 2004). ERSP was calculated from 1 Hz to the Nyquist frequency (500 Hz) with 1 Hz resolution with ripple centers at *t*=0 by computing and averaging fast Fourier transforms with Hanning window tapering. Each 1 Hz bin of the time-frequency matrix was normalized with respect to the mean power at -2000 to -1500 ms and masked with two-tailed bootstrapped significance (*N*=200) with false discovery rate (FDR) correction and *α*=0.05 using -2000 to -1500 ms as baseline. Grand average time-frequency plots were generated by averaging the average time-frequency plots of all channels for a given region (i.e., neocortex or hippocampus) and state (i.e., NREM or waking).

### Sleep and waking epoch selection

Epochs included in the study did not fall within at least 1 hour of a seizure and were not contaminated with frequent interictal spikes or artifacts. NREM periods were selected from continuous overnight recordings where the delta (0.5-2 Hz) analytic amplitude from the cortical channels was persistently increased (Table 1). Sleep epochs were confirmed by visual inspection to have normal appearing downstates, upstates, and spindles. Downstates, upstates, and spindles were also automatically detected, and quantification of these events showed they had the densities, amplitudes, and frequencies that are characteristic of NREM (see details in ‘Detection of downstates, upstates, and sleep spindles’ below). Waking periods were selected from continuous daytime recordings that had persistently low cortical delta as well as high cortical alpha (8-12 Hz), beta (20-40 Hz), and high gamma (70-190 Hz) analytic amplitudes. When the data included electrooculography (EOG; *N*=15/17 SEEG patients), waking periods also required that the 0.5-40 Hz analytic amplitude of the EOG trace was increased. Waking epochs were required to be separated from periods of increased delta analytic amplitude by at least 30 minutes.

### Ripple detection

The median center frequency of hippocampal ripples in 9 studies was ∼85 Hz in humans (Bragin et al., 1999; Staba et al., 2004; Clemens et al., 2007; Axmacher et al., 2008; Le Van Quyen et al., 2008; Staresina et al., 2015; Jiang et al., 2019b; Norman et al., 2019; Vaz et al., 2019), whereas in rodents, sharpwave-ripple frequency is ∼120-140 Hz (Buzsaki, 2015). In humans, putative 90 Hz hippocampal ripples have the same characteristic relation to sharpwaves, intra-hippocampal localization, modulation by sleep stage, and relation to cortical sleep waves as in rodents, and occur in hippocampi that have no signs of epileptic involvement. Furthermore, in rodent hippocampus the distinction between higher frequency gamma bursts and ripples is not always sharp, leading to the suggestion that for simplicity they both be referred to as ‘ripples’ (Stark et al., 2014), which we follow here.

Ripple detection was performed in the same way for all structures and states and was based on a previously described hippocampal ripple detection method (Jiang et al., 2019a; Jiang et al., 2019b). Requirements for inclusion and criteria for rejection were determined using an iterative process across patients, structures, and states. Data were first bandpassed at 60-120 Hz (6^th^ order Butterworth filter with zero-phase shift) and the top 20% of 20 ms moving root-mean-squared peaks were detected. It was further required that the maximum z-score of the analytic amplitude of the 70-100 Hz bandpass was greater than 3 and that there were at least 3 distinct oscillation cycles in the 120 Hz lowpassed signal, determined by shifting a 40 ms window in increments of 5 ms across ±50 ms relative to the ripple midpoint and requiring that at least 1 window have at least 3 peaks. Adjacent ripples within 25 ms were merged. Ripple centers were determined as the time of the maximum positive peak in the 70-100 Hz bandpass. Ripple onsets and offsets were identified on each side of the center peak when the 70-100 Hz analytic amplitude fell below 0.75 standard deviations above the mean. To reject epileptiform activities or artifacts, ripples were excluded if the absolute value of the z-score of the 100 Hz highpass exceeded 7 or if they occurred within 2 s of a ≥3mV/ms change in the broadband LFP. Ripples were also excluded if they fell within ±500 ms of putative interictal spikes, detected as described below. To exclude events that could be coupled across channels due to epileptiform activity, we excluded events that coincided with a putative interictal spike on any cortical or hippocampal SEEG channel included in the analyses. Events in SEEG recordings that had only one prominent cycle or deflection in the broadband LFP that manifested as multiple cycles above detection threshold in the 70-100 Hz bandpass were excluded if the largest valley-to-peak or peak-to-valley absolute amplitude in the broadband LFP was 2.5 times greater than the third largest. For each channel, the mean ripple-locked LFP and mean ripple band were visually examined to confirm that there were multiple prominent cycles at ripple frequency (70-100 Hz) and the mean time-frequency plot was examined to confirm there was a distinct increase in power within the 70-100 Hz band. In addition, multiple individual ripples in the broadband LFP and 70-100 Hz bandpass from each channel were visually examined to confirm that there were multiple cycles at ripple frequency without contamination by artifacts or epileptiform activity. Channels that did not contain ripples that met these criteria were excluded from the study. Of note, a recent study has identified ripples based on these criteria in a patient without epilepsy (Rubin et al., 2022).

### Subtraction of unit spikes from local field potentials

One challenge in detecting high frequency oscillations such as ripples in the LFP recorded by a microelectrode, is that unit spikes may ‘bleed through’ into the micro-LFP (Ray, 2015), thus resulting in the detection of spurious relationships between ripples and unit spikes. Although unit spikes are fast events, simply downsampling (which requires first low-passing to prevent aliasing) may not completely remove the influence of the action potential on the micro-LFP. To address this, we employed a modified unit spike waveform subtraction technique (Pesaran et al., 2002). Specifically, the average spike waveform (−500 to +1600 µs around the trough) of each unit was subtracted from the unfiltered 30 kHz Utah Array micro-LFP of the same channel centered on each spike. The data were then downsampled to 1 kHz and ripple detection was performed as described above. We confirmed that this method led to the detection of true oscillations through extensive visual confirmation of events in the 30 kHz micro-LFPs.

### Interictal spike detection and rejection

Ripples and other sleep waves were excluded if they were within ±500 ms from putative IIS detected as follows: A high frequency score was computed by smoothing the 70-190 Hz analytic amplitude with a 20 ms boxcar function and a spike template score was generated by computing the cross-covariance with a template interictal spike. The high frequency score was weighted by 13 and the spike score was weighted by 25, and an IIS was detected when these weighted sums exceeded 130. In each patient, detected IIS and intervening epochs were visually examined from hippocampal and cortical channels (when present) to confirm high detection sensitivity and specificity.

### Detection of downstates, upstates, and sleep spindles

Downstates and upstates were detected as previously described (Jiang et al., 2019a; Jiang et al., 2019b), where the broadband LFP from each channel was bandpassed from 0.1-4 Hz (6^th^ order Butterworth filter with zero-phase shift) and consecutive zero crossings separated by 0.25-3 s were selected. The top 10% amplitude peaks were selected and the polarity of each signal was inverted if needed so that downstates were negative and upstates were positive, by ensuring that the average analytic amplitude of the 70-190 Hz bandpass within ±100 ms around the peaks was greater for upstates than downstates. A total of 2,649,563 downstates were detected, with an average and standard deviation (across channels) density (occurrence rate) of 12.9±4.7 min^-1^ and amplitude of -237.3±169.8 µV. A total of 2,922,211 upstates were detected with a density of 14.6±4.7 min^-1^ and amplitude of 163.8±84.9 µV.

Spindles were detected as previously described (Hagler et al., 2018), where data were bandpassed at 10-16 Hz, then the absolute values were smoothed via convolution with a tapered 300 ms Tukey window, and median values were subtracted from each channel. Data were normalized by the median absolute deviation and spindles were detected when peaks exceeded 1 for at least 400 ms. Onsets and offsets were marked when these amplitudes fell below 1. Putative spindles that coincided with large increases in lower (4-8 Hz) or higher (18-25 Hz) band power were rejected to exclude broadband events as well as theta bursts, which may extend into the lower end of the spindle range (Gonzalez et al., 2018). A total of 694,168 spindles were detected with an average and standard deviation (across channels) density of 3.0±3.0 min^-1^, amplitude of 29.1±13.5 µV, oscillation frequency of 12.4±0.7 Hz, and duration of 633.5±67.2 ms.

### Ripple temporal relationships

Peri-cortical ripple time histograms of cortical sleep waves on the same channel were computed. Event counts were found in 50 ms bins for sleep waves within ±1500 ms around cortical ripple centers at *t*=0. A null distribution was generated by shuffling the event times relative to the ripples at *t*=0 within this 3 s window 200 times each. Pre-FDR *p*-values were computed by comparing the observed and null distributions within each bin over ±1000 ms for cortical sleep waves. These *p*-values were then FDR-corrected for the number of channels across patients multiplied by the number of bins per channel (Benjamini and Hochberg, 1995). A channel was considered to have a significant modulation if there were 3 or more consecutive bins with FDR-corrected *p*-values less than α=0.05. Whether events were leading or lagging cortical ripples at *t*=0 was computed for each channel with a two-sided binomial test with expected value of 0.5, using event counts in the 1000 ms before vs. 1000 ms after *t*=0 for sleep waves. Plots had 50 ms Gaussian smoothed event counts with 50 ms bins.

Conditional probabilities of a ripple given the following sleep waves or their sequences were computed: downstate, spindle, upstate, downstate–spindle, and spindle–upstate. A ripple given a spindle (R | SS) was determined if the ripple center occurred during the spindle (average spindle duration was 634 ms). A ripple was considered to precede an upstate (R | US) or follow a downstate (R | DS) if the ripple center occurred 634 ms before or after the peak of the upstate or downstate, respectively.

### Unit detection, classification, quality, and isolation

Unit detection and classification was performed according to our published procedures (Peyrache et al., 2012; Chan et al., 2014; Dehghani et al., 2016; Le Van Quyen et al., 2016; Teleńczuk et al., 2017; Eichenlaub et al., 2020; Dickey et al., 2021). Data were bandpassed at 300-3000 Hz and putative unit spikes were detected when the filtered signal exceeded 5 times the estimated standard deviation of the background noise. Units were k-means clustered using the first three principal components of each spike. Overlaid spikes were examined visually and those with abnormal waveforms were excluded. Based on their waveforms, firing rates, and autocorrelograms, action potentials were clustered as arising from putative pyramidal cells (PY) or interneurons (IN). PY had spike rates of ∼0.1-0.8 Hz, long valley-to-peak and half width intervals, sharp autocorrelations, and bimodal inter-spike interval (ISI) distributions, reflecting a propensity to fire in bursts. By contrast, IN had spike rates of ∼1-5 Hz, short valley-to-peak and half width intervals, broad autocorrelations, and a predominantly unimodal ISI distribution. The Utah Array patient included in the study had 69 PY with a total of 231,922 spikes, as well as 23 IN with a total of 462,246 spikes. The average and standard deviation PY valley-to-peak amplitude was 44.8±12.9 µV, spike rate was 0.28±0.17 Hz, valley-to-peak width was 0.49±0.5 ms, half-peak width was 0.62±0.04 ms, and bursting index was 0.03±0.02. The average and standard deviation IN valley-to-peak amplitude was 28.9±11.9 µV, spike rate was 1.67±1.63 Hz, valley-to-peak width was 0.29±0.05 ms, half-peak width was 0.34±0.05 ms, and bursting index was 0.01±0.01. Valley-to-peak amplitude, spike rate, valley-to-peak width, half-peak width, and bursting index were all significantly different between PY and IN (*p*<0.0001, two-sided two-sample *t*-test).

Single unit quality and isolation were confirmed according to previously established guidelines (Kamiński et al., 2020). Unit spikes were verified to well-exceed the noise floor based on large peak signal-to-noise ratios (PY: 10.1±3.4; IN: 5.9±3.1). Since the neuronal spiking refractory period is about 3 ms, the percent of ISIs less than 3 ms estimates the degree of single unit contamination by spikes from different units, which was very low among the units included in this study (PY: 0.12±0.15%; IN: 0.31±0.50%). Furthermore, single units detected on the same contact were highly separable according to their projection distances (Pouzat et al., 2002) (PY: 49.2±25.8 SD; IN: 50.9±28.6 SD). Lastly, temporal stability of unit spikes over time was confirmed based on consistency of the mean waveform shape and amplitude of each unit across recording quartiles.

### Analyses of unit spiking during ripples

Unit spiking was analyzed with respect to local ripples detected on the same contact. Ripple phases of unit spikes were determined by finding the angle of the Hilbert transform of the 70-100 Hz bandpassed signal (zero-phase shift) at the times of the spikes. The circular mean ripple phase was determined for each unit according to Berens (2009). Phase analyses were only performed on units that had at least 30 spikes during local ripples. The Ralyleigh test was used to assess for unimodal deviation from uniformity of the circular mean phases across units. PY-PY and PY-IN unit pair co-firing within 5 ms was assessed based on Dickey et al. (2021). To evaluate co-firing when there was a ripple at either or both sites vs. baseline when there were no ripples occurring, we randomly selected non-ripple epochs matched in number and duration to the ripples but during which there were no ripples detected on either channel. Co-firing during ripples with observed vs. random spike times was computed by shuffling the spike times of each unit within the intervals of the ripples.

### Experimental design and statistical analyses

All statistical tests were evaluated with α=0.05. All *p*-values involving multiple comparisons were FDR-corrected according to Benjamini and Hochberg (1995). FDR corrections across channel pairs were done across all channels pairs from all patients included in the analysis. Box-and-whisker plots show median, mean, and interquartile range, with whiskers indicating 1.5 × interquartile range with outliers omitted. Kernel density plots were produced using methods from Bechtold (2016). Significance and shuffling statistics of peri-ripple time histograms were computed as described above. Unit spiking statistics were computed as described above.

Statistical comparisons between ripple characteristics were performed using two-sided paired or two-sided two-sample *t*-tests between channel means. Differences in ripple characteristics between states (NREM or waking) or regions (cortex or hippocampus) were determined using linear mixed-effects models with state and region as fixed effects and patient as a random effect, according to the following model:

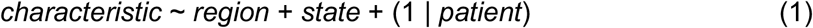

Factors that had *p*<0.05 then underwent *post*-*hoc* testing with the following models. These *p*-values were then FDR-corrected for multiple comparisons.

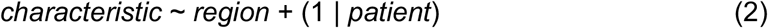

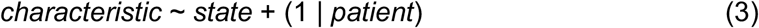

Significance of linear correlations (i.e., ripple characteristics vs. myelination index or hippocampal connectivity density) were assessed when there were at least 10 data points using linear mixed-effects models with patient as random effect, according to the following model:

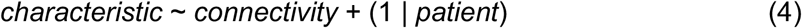

## Acknowledgements

We thank Adam Niese, Christine Smith, Christopher Gonzalez, Daniel Cleary, Eran Mukamel, Erik Kaestner, Jacob Garrett, Maxim Bazhenov, Terrence Sejnowski, and Zarek Siegel for their support. This work was supported by NIMH (1RF1MH117155-01, T32 MH020002) and ONR-MURI (N00014-16-1-2829).

## Data availability

The de-identified raw data that support the findings of this study are available from the corresponding authors upon reasonable request provided that the data sharing agreement and patient consent permit that sharing.

## Code availability

The code that support the findings of this study are available from the corresponding authors upon reasonable request.

