## Supplementary Information for "Cortical ripples during NREM sleep and waking in humans"

| Cortical Region | Desikan Parcels |
|---|---|
| Orbitofrontal | frontal pole, lateral orbitofrontal, medial orbitofrontal, pars orbitalis |
| Prefrontal | caudal middle frontal, pars opercularis, pars triangularis, rostral middle frontal, superior frontal |
| Rolandic | paracentral, postcentral, precentral |
| Parietal | inferior parietal, superior parietal, supramarginal, precuneus |
| Occipital | cuneus, pericalcarine, lateral occipital |
| Superior Temporal / Insula | insula, superior temporal, transverse temporal |
| Lateral Temporal | banks of superior temporal gyrus, inferior temporal, middle temporal, |
| Ventral Temporal | entorhinal, fusiform, lingual, parahippocampal, temporal pole, isthmus cingulate |
| Cingulate | caudal anterior cingulate, rostral anterior cingulate, posterior cingulate |

**Supplementary Table 2-1. Relationships between cortical regions reported and Desikan parcels.** First column lists the cortical regions described in this study, which are derived from an amalgamation of the cortical parcels from Desikan et al. (2006) listed in the second column.

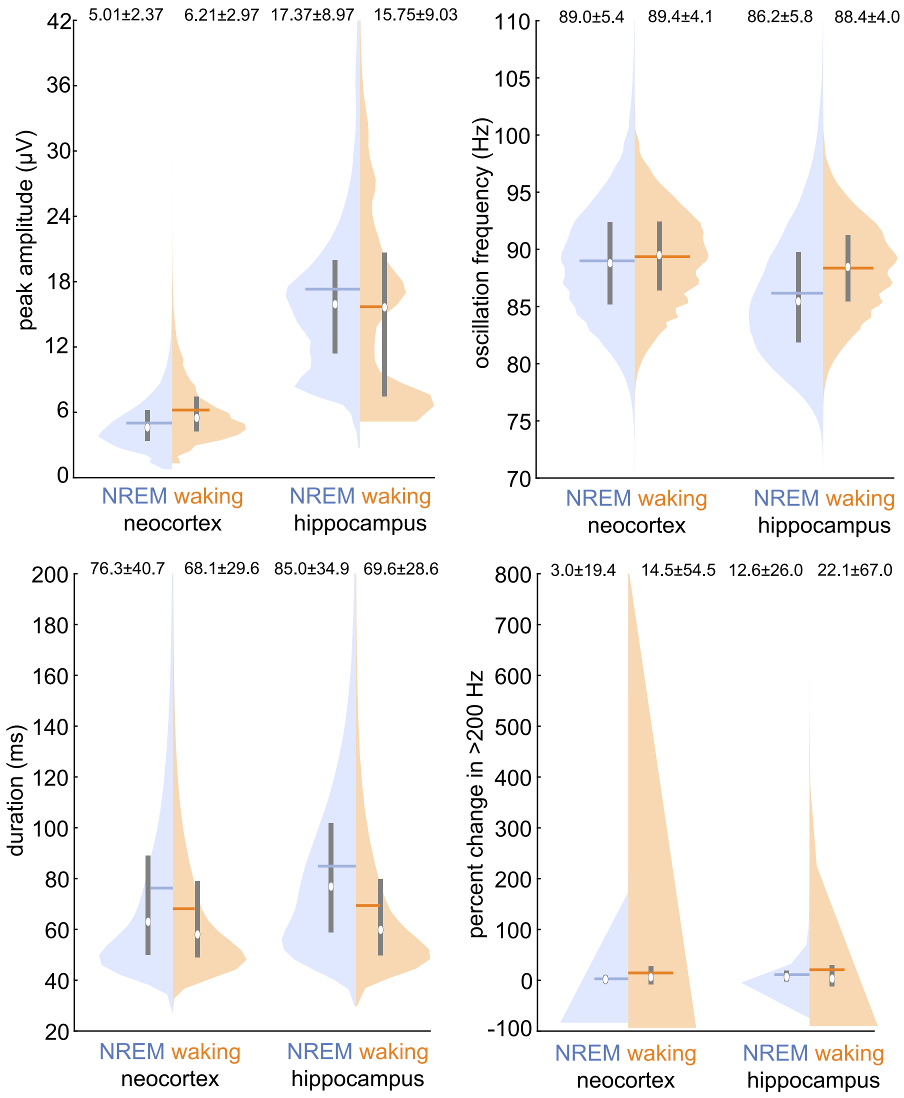


**Supplementary Figure 3-1. Characteristics of individual cortical ripples.** Histograms of ripple characteristics across individual events ($N_{NREM}$=1,906,502, $N_{waking}$=3,415,232) from all channels (*N*=273 cortical, *N*=28 hippocampal) from all SEEG patients (S1-17). Values are means and standard deviations. Circles show medians; horizontal lines, means; vertical lines, interquartile ranges.

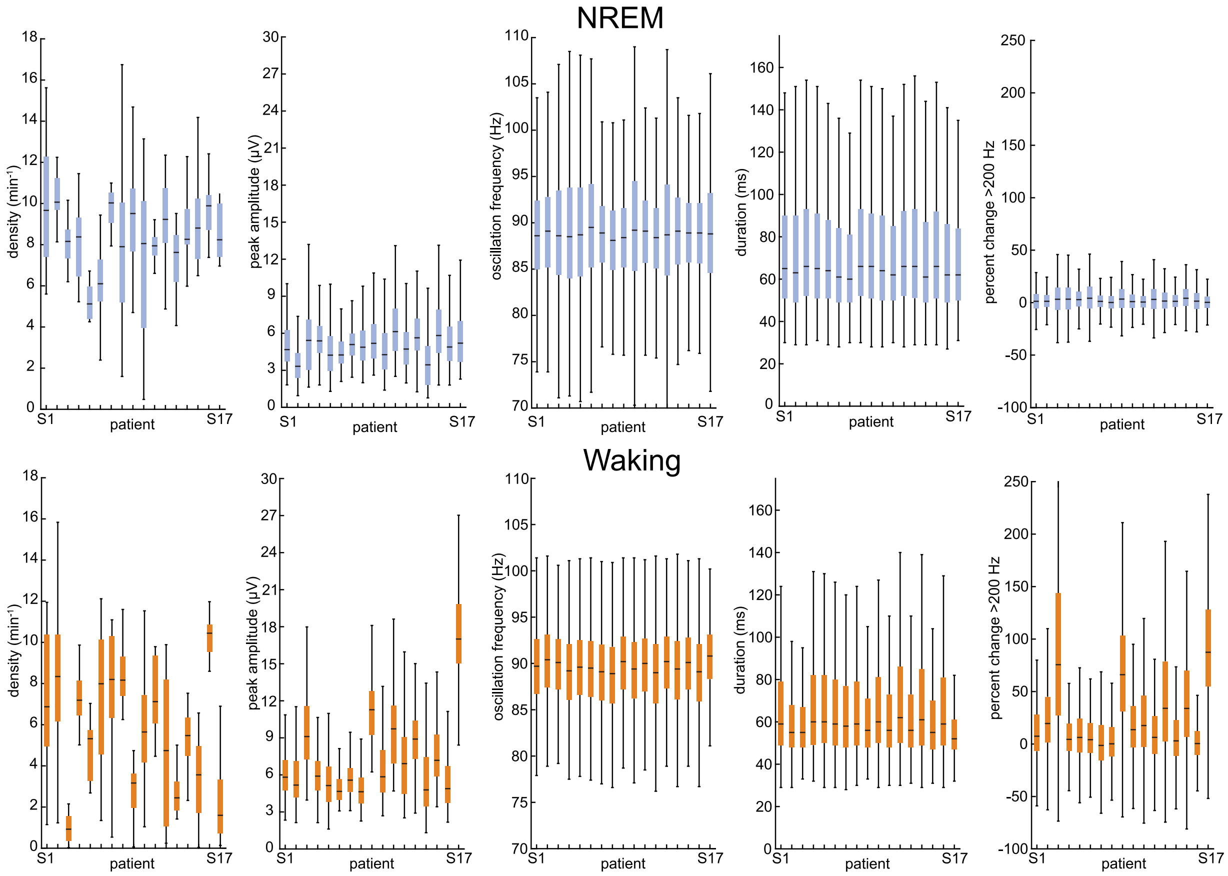


**Supplementary Figure 3-2. Cortical ripple characteristics by patient.** Cortical ripple characteristics are consistent across patients (S1-17). Horizontal lines show medians; boxes, interquartile ranges; whiskers, 1.5 × interquartile range.

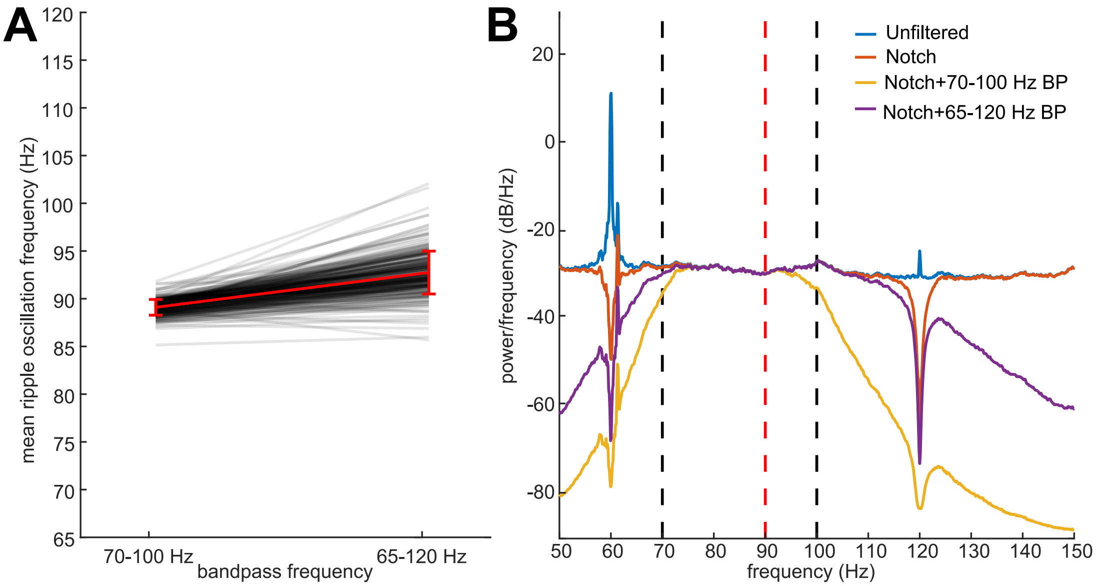


**Supplementary Figure 3-3. The ~90 Hz ripple oscillation frequency is not explained by filtering or detection bandpass.** (**A**) Mean and standard deviation NREM ripple oscillation frequency (red line) across channels ($N$=273; black lines) from all SEEG patients (S1-17) is highly similar when using either a 70-100 Hz bandpass (mean±SD=89.1±0.8 Hz) or a 65-120 Hz bandpass (92.7±2.2 Hz). (**B**) Average power spectral density of unfiltered, notched (60 Hz and harmonics), notched and 70-100 Hz bandpassed, as well as notched and 65-120 Hz bandpassed ($N$=1000 randomly selected 27 s long epochs over 450 min of recording from patient S2). Dashed black lines indicate 70-100 Hz bandpass used for the main analyses in this study. Dashed red line shows 90 Hz, the approximate average ripple frequency across states, structures, and filter settings.

| Graphoelement \| Ripple | Signif. Modulation | Signif. Sidedness | Cort-R Leading |
|---|---|---|---|
| Downstate \| Cort-R | 94.51% (258/273) | 63.57% (164/258) | 33.54% (56/164 |
| Spindle \| Cort-R | 29.3% (80/273) | 73.75% (59/80) | 28.81% (17/59) |
| Upstate \| Cort-R | 95.24% (260/273) | 83.85% (218/260) | 86.70% (189/218) |

**Supplementary Figure 8-1. Sleep wave–ripple coupling.** Proportion of channels with significant peri-ripple modulations of sleep waves detected on the same channels within ±1000 ms (e.g., Upstate | Cort-R represents upstate peaks relative to cortical ripples at $t$=0; one-sided randomization test, 200 shuffles, 50 ms non-overlapping bins, 2 consecutive bins with $p<0.05$ required for significance), and those with significant modulations that had significant sidedness preference around $t$=0 ($p<0.05$, one-sided binomial test, -1000 to -1 ms vs. 1 to 1000 ms, expected=0.5), and those with significant sidedness around 0 that had cortical ripples leading sleep waves (according to counts in -1000 to -1 ms vs. 1 to 1000 ms). In the calculations, upstate and downstate times were peaks and spindle times were onsets. *P*-values were FDR-corrected across channels and bins. See Figure 8A for single sweep example, Figure 8B-D for peri-ripple time histograms of cortical sleep waves, and Figure 8E-G for graphical representations of conditional probabilities.
